# Pre-existing levels of pro-survival proteins and induction of BCL-XL dictate cell fate after p53 activation

**DOI:** 10.64898/2026.07.01.735749

**Authors:** Allan Shuai Huang, Elizabeth Lieschke, Pedro L Baldoni, Annabella F Thomas, Julia M Marchingo, Lauren Whelan, Grace Khuu, Eddie La Marca, Michael Milevskiy, Aisling M Ross, Tim Johanson, Maggie Potts, Leonie Gibson, Vineet Vaibhav, Laura Dagley, Amar Balihodcik, Michael Dengler, Ziyan Liu, Kaiming Li, Gordon K Smyth, Gemma Kelly, Andreas Strasser

**Affiliations:** The Walter and Eliza Hall Institute (WEHI), Melbourne, Australia; Department of Medical Biology, The University of Melbourne, Melbourne, Australia; University of Graz, Graz, Austria

**Author notes:** Address for correspondence: Gemma Kelly or Andreas Strasser, The Walter and Eliza Hall Institute (WEHI), 1G Royal Parade, Parkville, Victoria 3052, Australia. ASH, EL and PLB share first authorship. GKS, GK and AS share senior authorship.

## Abstract

TP53 (also called TRP53 or p53) is a critical tumour suppressor that prevents cancer development by inducing a transcriptional program which can lead to diverse cellular responses, most prominently, cell proliferation arrest/senescence with survival of cells or cell death by apoptosis. Why distinct cell types undergo different outcomes after p53 activation remains unclear. Using integrated RNA-sequencing, proteomic and functional analyses across a diverse range of murine primary cell types, we demonstrate that cell fate is governed by the balance between pro-survival BCL-2 and pro-apoptotic BH3-only proteins. Cells resistant to apoptosis displays a higher starting ratio of pro-survival BCL-2 to pro-apoptotic BH3-only proteins, along with transcriptional upregulation of the pro-survival gene *Bcl2l1*, encoding BCL-XL. This control of cell fate is also seen in human wild-type p53 cancer cell lines. These findings reveal the mechanism for understanding p53-driven cell fate decisions, suggest therapeutic strategies to shift p53-induced cell proliferation arrest/senescence toward apoptotic cell death and allowed generation of an RNAseq data-based predictor of outcome for cancer cells after p53 activation.

## Introduction

TP53 (also called TRP53 in mice or p53 in general) is a critical tumour suppressor and transcription factor that safeguards cellular integrity by preventing malignant transformation (Lane 1992, Bieging, Mello et al. 2014, Kastenhuber and Lowe 2017). Upon activation in response to diverse stresses, such as oncogene expression, DNA damage or nutrient deprivation, p53 induces a transcriptional program by directly activating ∼500 target genes, and several more indirectly (Kenzelmann Broz, Spano Mello et al. 2013, Thomas, Kelly et al. 2022). This unleashes transcriptional programs for induction of cell cycle arrest/cell senescence, apoptotic cell death, DNA damage repair and adaptation of cellular metabolism. How after activation of p53 cells make the fundamental decision between cell cycle arrest/senescence with survival versus apoptotic cell death remains incompletely understood.

It has been postulated that cell survival with proliferation arrest (and senescence) versus cell death after p53 activation is determined by selective transcriptional induction of either of the direct p53 target genes *p21(Cdkn1a)* encoding a cyclin dependent kinas inhibitor (CDKI) critical for G1/S boundary cell cycle arrest/senescence (Deng, Zhang et al. 1995), or *Puma(Bbc3)* and *Noxa(Pmaip1)* encoding initiators of apoptosis (Nakano and Vousden 2001, Yu, Zhang et al. 2001, Jeffers, Parganas et al. 2003, Villunger, Michalak et al. 2003). However, some RNAseq and qRT-PCR analysis revealed that both *p21(Cdkn1a)* as well as the pro-apoptotic genes *Puma(Bbc3)* and *Noxa(Pmaip1)* were induced in both cell types that survive and undergo cell cycle arrest/senescence following p53 activation as well as in ones that undergo apoptosis (Kenzelmann Broz, Spano Mello et al. 2013, Valente, Aubrey et al. 2016, Moyer, Wasylishen et al. 2020). These findings challenge the idea that differential transcriptional activation of *p21(Cdkn1a)* or *Pum(Bbc3)* and *Noxa(Pmaip1)* alone determines cell fate, indicating that additional factors must govern the survival versus death decision after p53 activation.

Understanding why cells undergo one cellular outcome but not the other has implications for cancer therapy. Although both cell cycle arrest/senescence and apoptosis suppress tumour growth, proliferation arrest, even with cell senescence, represents a potentially reversible state. Arrested cells can escape and re-enter the cell cycle (Waldman, Zhang et al. 1997), thereby contributing to tumour relapse. Moreover, senescent cells may promote a pro-inflammatory microenvironment through senescence-associated secreted proteins (SASP) that can promote tumour expansion (Wang, Han et al. 2024, Cao, Li et al. 2025). In contract, apoptotic cell death permanently eliminates cancer cells and therefore is a therapeutically more desirable outcome.

To explore the mechanisms governing cell fate after p53 activation, we performed RNA-sequencing and proteomic analysis across a diverse range of murine primary cell types. This revealed a much higher ratio of anti-apoptotic BCL-2 proteins to pro-apoptotic BH3-only proteins, along with further transcriptional upregulation of the pro-survival gene *Bcl2l1* (encodes BCL-XL) in cells that survive after p53 activation compared with cells that undergo apoptosis. Enforced increases in BCL-XL (or MCL-1) allowed cells to survive that would normally die after p53 activation. Conversely, blunting the pro-survival function of BCL-XL (and/or MCL-1) killed cell types that typically survive after p53 activation. Using this discovery, we devised a model based on the ratio of the basal levels of mRNAs for pro-survival BCL-2 versus pro-apoptotic BH3-only proteins that predicts faithfully whether upon p53 activation human cancer cells will survive and undergo cell cycle arrest or die by apoptosis. Together, our findings establish a mechanistic and predictive framework for understanding, and potentially manipulating, the cell survival-versus-death decision following p53 activation.

## Results

### p53 activation induces distinct cellular outcomes across different primary murine cell types

Activation of p53 can induce diverse cellular responses, including cell cycle arrest/senescence with cell survival versus apoptotic cell death. To determine how different primary murine cell types respond to p53 activation, we treated murine dermal fibroblasts (MDFs), bone marrow–derived macrophages (BMDMs), activated T cells, activated B cells, thymocytes, and pre-B cells with the MDM2 inhibitor nutlin-3a that activates p53 in a non-genotoxic manner (Vassilev, Vu et al. 2004) (Figure 1A). Apoptosis and cell cycle distribution were assessed by flow cytometry. Following p53 activation, MDFs and BMDMs were largely resistant to apoptosis. After 72 h of nutlin-3a treatment, more than 80% of cells remained viable (Figure 1B). Instead, these cells underwent cell cycle arrest, as evidenced by their accumulation in G0/G1 and G2/M phases and a corresponding reduction in cells in S phase (Figure 1C, D). In contrast, thymocytes and pre-B cells were highly sensitive to p53 activation induced apoptosis, with nearly 100% of cells having died within 24 h of treatment with nutlin-3a (Figure 1B). Activated T and B cells exhibited an intermediate cellular outcome. Approximately 50% of cells underwent apoptosis within 24 h of treatment with nutlin-3a (Figure 1B), while the remaining viable cells had undergone cell cycle arrest, evidenced by accumulation in the G0/G1 phase and reduction of cells in S phase (Figure 1E and 1F).

**Figure 1:**
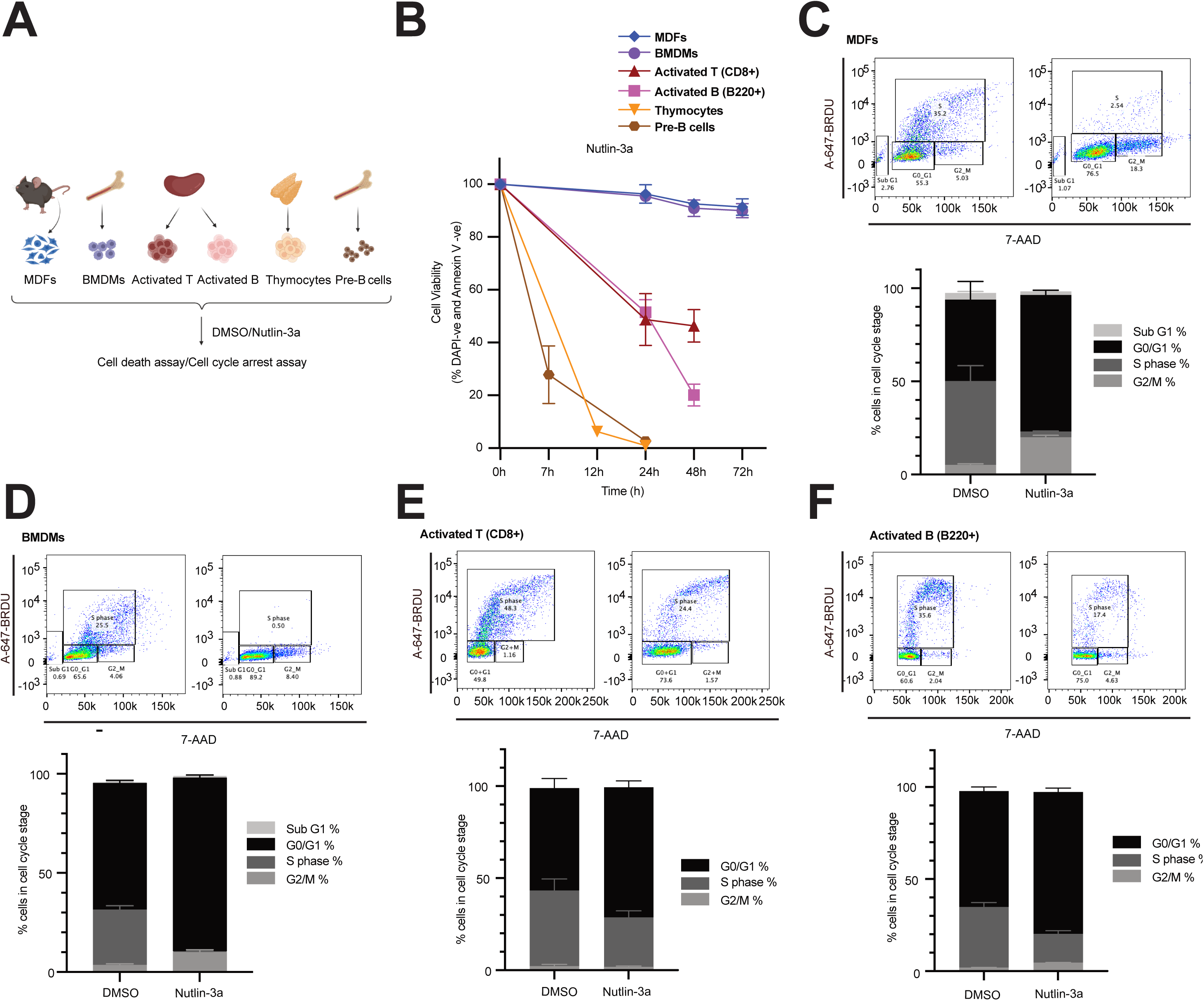
p53 activation induces distinct cellular outcomes across a range of primary murine cell types. **A**: Schematic of the primary murine cell types derived from C57BL/6 mice used in this study. **B**: MDFs, BMDMs, activated T cells, activated B cells, thymocytes, and pre-B cells from C57BL/6 mice were treated with 10 µM nutlin-3a for the indicated time points (MDFs and BMDMs were analysed after 24 h, 48 h and 72 h, activated T cells and activated B cells were analysed after 24 h and 48 h, thymocytes were analysed after 12 h and 24 h, pre-B cells were analysed after 7 h and 24 h). Apoptosis was assessed by Annexin-V plus DAPI staining followed by flow cytometry (Annexin V-DAPI- cells were deemed live cells). Data are presented as mean ± SD (n = 3 biological replicates). **C–F**: Cell cycle distribution of MDFs (C), BMDMs (D), activated T cells (E) and activated B cells (F) after 24 h treatment with 10 µM nutlin-3a with cells pulsed with BrdU for the last 4h. Cell cycle distribution was determined by staining for BrdU incorporation and DNA content (7-AAD) and cells were divided into the following stages of the cell cycle: sub-G1 (apoptotic), G0/G1, S, or G2/M. For activated T cells and B cells, dead cells were excluded, and cell cycle analysis was performed on viable cells only. Data are presented as mean ± SD (n = 3 biological replicates).

### *p21(Cdkn1a)* and *Puma(Bbc3)* are induced concomitantly by p53 within a single cell

To determine if selective induction of the direct p53 target genes *p21(Cdkn1a)* or *Puma(Bbc3)* explains the distinct cellular outcomes across different primary murine cell types, their mRNA levels were examined at several time points of treatment with nutlin-3a (2, 4, 6, 8 and 24 h) by qRT-PCR. In MDFs, which predominantly undergo cell cycle arrest upon p53 activation, both *p21(Cdkn1a)* and *Puma(Bbc3)* transcripts were robustly induced compared with vehicle (DMSO) treated control cells (Figure 2A, B). No induction of these genes was observed in MDFs from *p53^−/−^* mice, confirming that nutlin-3a strictly acts in a p53-dependent manner as reported (Vassilev, Vu et al. 2004). In activated T cells, which displayed mixed cellular outcomes, and in thymocytes, which are highly sensitive to p53-induced apoptosis, both *p21(Cdkn1a)* and *Puma(Bbc3)* were upregulated upon treatment with nutlin-3a (Figure 2C-F). Even though this qRT-PCR analysis and published RNAseq data (Kenzelmann Broz, Spano Mello et al. 2013) revealed induction of both cell cycle arrest inducing (*p21*) as well as apoptosis initiating genes (*Puma*, *Noxa*) in the same cells, this was done at a population level. Thus, it has not been formally demonstrated that these genes that encode proteins operating in opposed cellular processes are actually induced within a single cell across cell types with different outcomes after p53 activation. To assess simultaneous induction of *p21* and *Puma* at the single-cell level, we generated double-reporter mice by intercrossing *p21-IRES-GFP* and *Puma-tdTomato* reporter strains (Lieschke, Thomas et al. 2024). In these mice, *p21* transcription is reported by GFP expression and *Puma* transcription by tdTomato fluorescence, enabling simultaneous monitoring induction of both target genes within individual cells. MDFs, BMDMs, activated T cells and activated B cells were treated with nutlin-3a, and reporter expression analysed by flow cytometry. Treatment with nutlin-3a treatment led to increased GFP and tdTomato fluorescence across all cell types examined, with a clear correlation between the intensity of the signals of the two reports; i.e. cells with high levels of GFP (*p21-IRES-GFP* reporter) also had high levels of tdTomato (*Puma-tdTomato* reporter) (Figure 2H and Supplementary Figure 1). This reveals transcriptional induction of both *p21* and *Puma* within single cells across cell types that display opposite outcomes after activation of p53.

**Figure 2:**
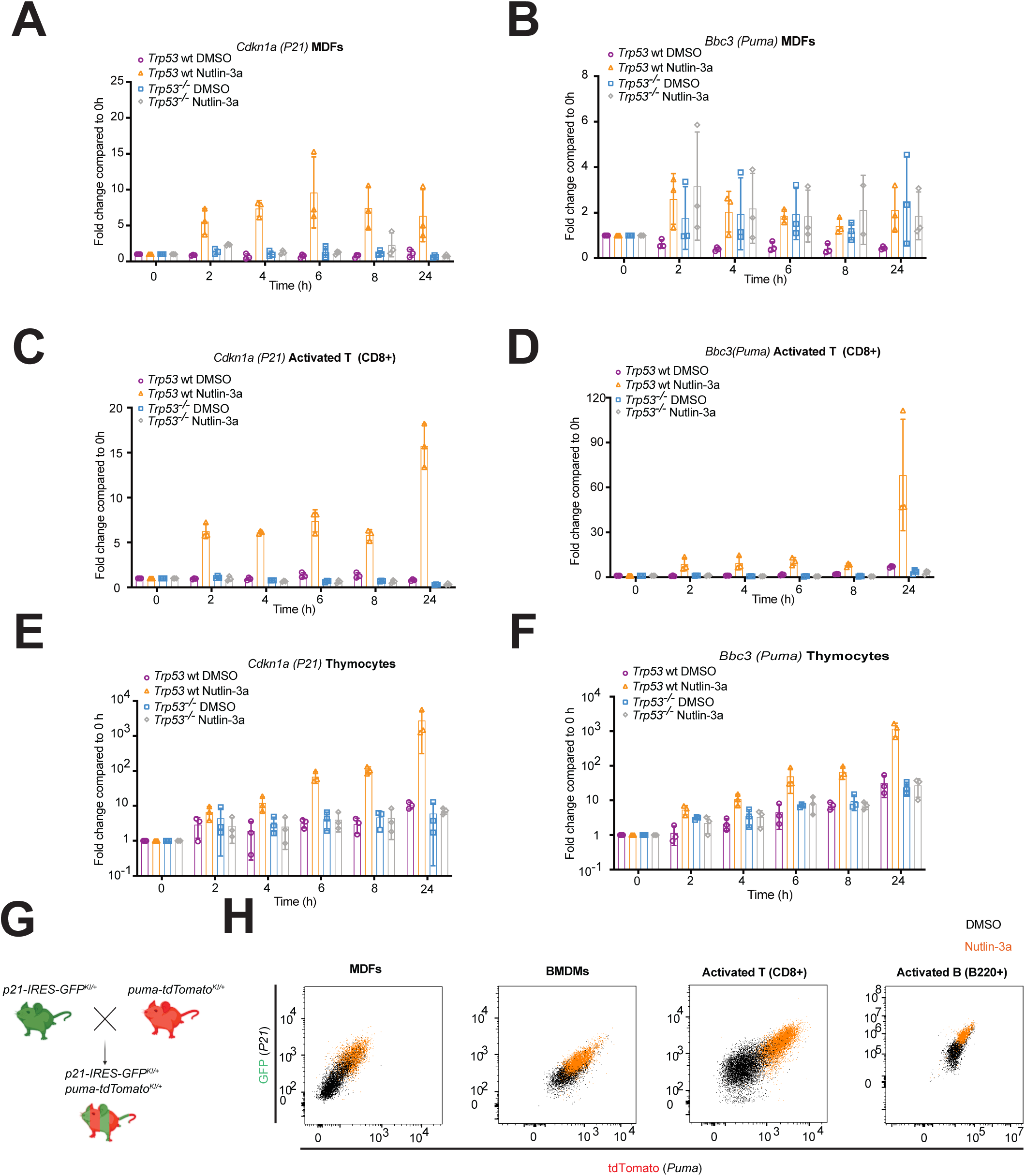
p53 activation causes an increase in both *P21*(*Cdkn1a)* and *Puma*(*Bbc3)* mRNA within a single cell regardless of whether it will survive and undergo proliferation arrest or undergo apoptotic death. **A-F**: RNA was extracted from MDFs, activated T cells and thymocytes from wt and *p53^-/-^* mice following treatment *in vitro* with DMSO (vehicle control) or 10 µM nutlin-3a for 2 h, 4 h, 6 h, 8 h or 24 h. qRT-PCR analysis was performed to measure the mRNA levels of the p53 target genes *P21*(*Cdkn1a)* and *Puma*(*Bbc3)*. Data were normalised using the ΔΔCT method with *Hmbs* used as a housekeeping gene. Data are presented as fold-change compared to DMSO treated samples. Data are presented as mean ± SD (n = 3 mice of each genotype and treatment). **G**: Generation of *p21-IRES-GFP^KI/+^*;*Puma-tdTomato^KI/+^*double reporter mice by crossing *p21-IRES-GFP^KI/+^* and *Puma-tdTomato^KI/+^*strains. **H**: Representative flow cytometry plots of MDFs, BMDMs, activated T cells and activated B cells from *p21-IRES-GFP^KI/+^;Puma-tdTomato^KI/+^*double reporter mice following 24 h treatment *in vitro* with DMSO (vehicle control) or 10 µM nutlin-3a, showing GFP (*p21*) and tdTomato (*puma*) reporter levels. For activated T cells and activated B cells, the caspase inhibitor Q-VD-OPh (QVD) was included to prevent cell demolition due to apoptosis. Data shown are representative of n=3 independent mice. Summary plots of GFP and tdTomato expression corresponding to panel H are shown in **Supplementary Figure 1**, presented as fold-change relative to the DMSO treated control cells at 0 h.

To examine whether p21 is required for the observed cell fate decisions, as proposed in some but not other studies (Bharti, Watkins et al. 2022, Bell, Blair et al. 2024), we compared cellular outcomes between wild-type and *p21 (Cdkn1a)^−/−^* cells across each primary cell type. No substantial differences were observed between cells of the different genotypes. Similar to their wild-type counterparts, p21-deficient MDFs and BMDMs did not undergo apoptosis after p53 activation. Both wild-type and *p21 (Cdkn1a)^−/−^* MDFs underwent cell cycle arrest, as evidenced by their accumulation in G0/G1 and G2/M phases and a corresponding reduction of cells in S phase; however, the extent of arrest was reduced in the absence of p21. In contrast, whereas wild-type BMDMs exhibited a clear p53-induced cell cycle arrest, loss of *p21(Cdkn1a)* completely abrogated this response Supplementary Figure 2A–D). After treatment with nutlin-3a p21-deficient mitogen activated T and B lymphocytes exhibited cellular outcomes comparable to wild-type cells, with ∼50% of cells undergoing apoptosis and the remaining population displaying cell cycle arrest, although the extent of cell cycle arrest was reduced or abrogated in p21-deficient cells compared to wild-type cells (Supplementary Figure 2E–H).

These findings demonstrate that p53 simultaneously induces transcription of both cell cycle arrest and apoptosis inducing target genes within a single cell, irrespective of whether it will survive and undergo cell cycle arrest/senescence or die by apoptosis, and that p21 is not required for determination of cell survival versus apoptotic death.

### BCL-XL upregulation at both the transcript and protein levels in cells that do not undergo apoptosis after activation of p53

Since differential induction of *p21 (Cdkn1a)* and *Puma (Bbc3*)as well as pro-apoptotic genes (*Noxa/Pmaip1*, *Bim/Bcl2l11* and *Bax)* could not explain cell fate after p53 activation, we performed a large-scale RNA sequencing analysis to systematically identify genes that are induced or repressed after treatment with nutlin-3a in six different murine primary cell types representing distinct cell fate responses (Figure 1A), using matched *p53^-/-^*cells as controls. Our aim was to identify genes were selectively induced only in cells undergoing apoptosis or, conversely, only in cells that survive and undergo cell cycle arrest after p53 activation.

Multidimensional scaling (MDS) analysis revealed that samples clustered primarily by cell type rather than treatment condition (Supplementary Figure 3A and B), indicating that p53 activation does not override intrinsic cellular identity. Pathway-focused interrogation of genes associated with KEGG categories including p53 signalling, apoptosis, cell cycle, and senescence confirmed global induction or repression of canonical p53 pathway components in all wild-type cell types (Figure 3A). As expected, no such changes in gene expression were detected in the cells from *p53^-/-^*mice after treatment with nutlin-3a (Figure 3A). To further interrogate pathway-level responses, we performed self-contained gene set testing using curated gene lists (Supplementary table 1) for canonical p53-associated pathways. As expected, p53 signalling was significantly activated across all cell types after treatment with nutlin-3a. Notably and consistent with the results from the analysis of the *p21-IRES-GFP*;*Puma-tdTomato* double-reporter cells (Figure 2H and Supplementary Figure 1), apoptosis- and senescence-associated gene sets were significantly upregulated in cell types that survived upon p53 activation but were also detectable in the cells destined for apoptotic death. In contrast, cell cycle–associated gene sets were selectively downregulated in cell types undergoing cell cycle arrest (MDFs, BMDMs, and activated T and B lymphocytes), but not in thymocytes or pre-B cells, which reside in the G0 cell cycle state and rapidly undergo apoptosis (Figure 3B and Supplementary Figure 4). Together, these findings indicate that cell fate following p53 activation cannot be explained by selective engagement of canonical pathways, as these programs - particularly apoptosis and senescence - are activated across cell types regardless of their fate.

**Figure 3.**
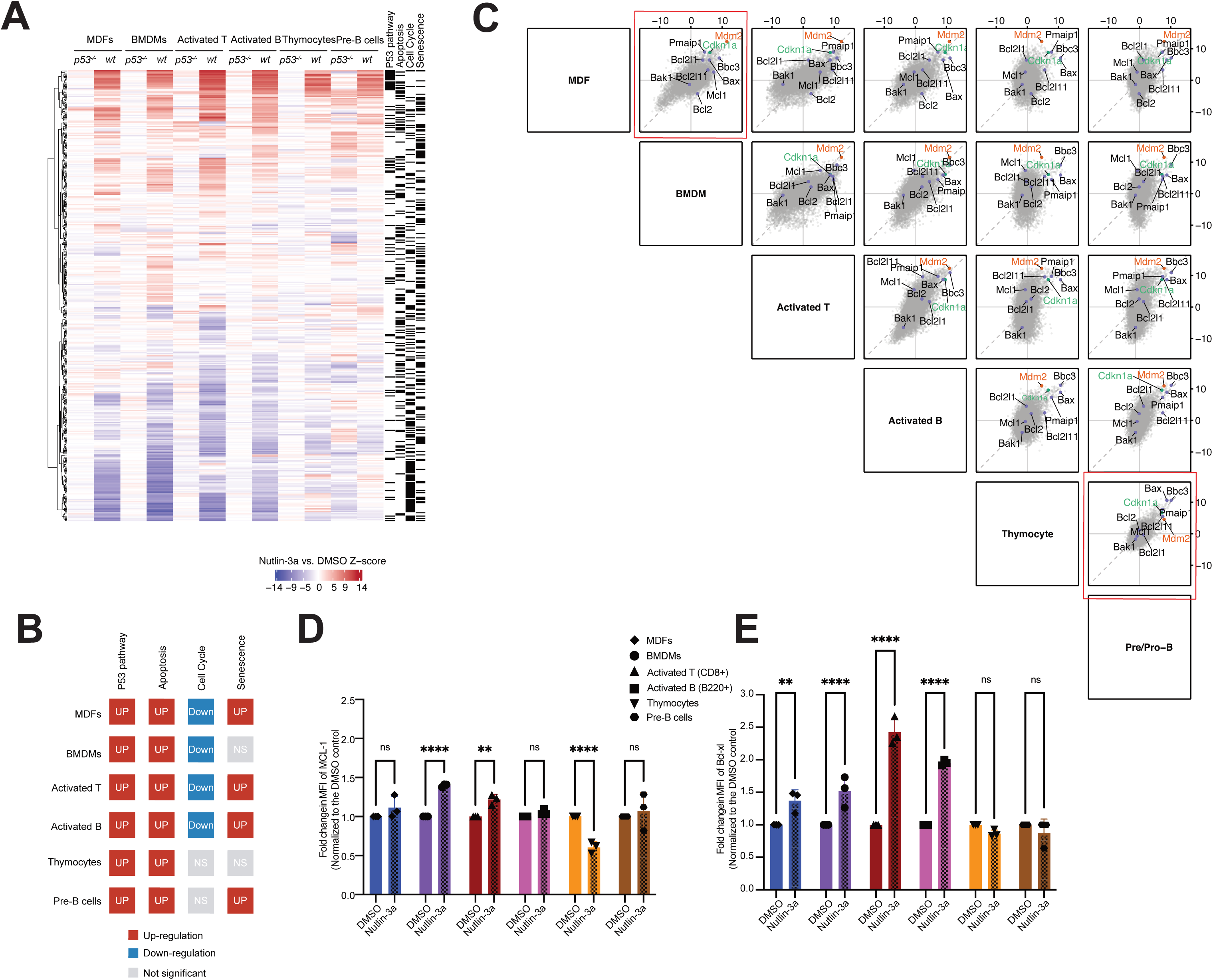
*Bcl2l1* encoding BCL-XL is upregulated at both transcript and protein levels in cells that survive and undergo proliferation arrest after p53 activation. **A:** Heatmap showing expression of genes from KEGG pathways related to p53 signalling, apoptosis, cell cycle, and senescence. Z-scores were calculated from t-statistics comparing 6 h nutlin-3a treated versus DMSO treated samples in the presence of 25 µM QVD-OPH within each cell type (MDFs, BMDMs, activated T cells, activated B cells, thymocytes, and pre-B cells from wild-type and *p53⁻/⁻* mice). **B:** Summary plot showing the direction of change from self-contained gene set testing for each pathway within each cell type. Individual barcode plots are shown in **Supplementary Figure 4.** **C:** Pairwise plots showing Z-scores derived from t-statistics comparing nutlin-3a treatment versus DMSO treatment within each wild-type cell type. **D** and **E:** Summary of intracellular flow cytometric analysis of MCL-1 (**D**) and BCL-XL (**E**) Protein levels in MDFs, BMDMs, activated T cells, activated B cells, thymocytes, and pre-B cells from wild-type mice following 24 h treatment with DMSO or 10 µM nutlin-3a in the presence of 25 µM QVD-OPH. Each symbol represents cells from an individual mouse (n = 3). Data are presented as mean ± SD. Protein abundance was quantified by median fluorescence intensity (MFI) following subtraction of the corresponding Ig isotype control staining intensity and normalisation to DMSO treated control cells. Statistical significance was determined using two-way ANOVA with Dunnett’s multiple comparisons test (P < 0.05; P < 0.01; P < 0.001; P < 0.0001; ns, not significant). Representative flow cytometry plots are shown in **Supplementary Figure 6**.

We next focused on individual key genes associated with cell fate regulation, including cell cycle (p21/*Cdkn1a)*, feedback regulation of p53 (*Mdm2*), and apoptosis-related genes, given their central roles in determining the cell survival versus apoptotic death decision. To identify shared and cell type–specific transcriptional responses, we performed pairwise comparisons of differentially expressed genes upon p53 activation across the six cell types. Consistent with our analysis of the cells from the *p21-IRES-GFP;Puma-tdTomato* double reporter mice (Figure 2), both *p21* (*Cdkn1a*) and *Puma* (*Bbc3*) were among the genes robustly induced across all examined cell types. In addition, the RNA-seq data revealed induction of other canonical p53 target genes, including those encoding the effector of apoptosis BAX, the BH3 only protein NOXA (*Pmaip1*), and the negative regulator of p53, MDM2 (Figure 3C). Strikingly differential patterns emerged when examining genes for pro-survival BCL-2 family members. Transcripts encoding BCL-XL (*Bcl2l1*) were significantly increased in nutlin-3a treated MDFs, BMDMs and activated T as well as activated B cells - cell types that resist apoptosis or exhibit mixed cellular outcomes - but not in thymocytes or pre-B cells, which undergo rapid apoptosis following p53 activation (Figure 3C). *Mcl-1* transcripts were significantly elevated in BMDMs and activated T cells, but this was not seen in any of the cell types that die rapidly after p53 activation (Figure 3C). In contrast, genes encoding pro-apoptotic BIM (*Bcl2l11*), BAX and NOXA (*Pmaip1*) were upregulated across all cell types examined, and those encoding the apoptosis effector BAK (*Bak1*) or pro-survival BCL-2 did not significantly change (Figure 3C). Importantly, none of these transcriptional alterations were seen in the corresponding cells from p53-deficient mice (Supplementary Figure 5). This validates that both the shared p53 transcriptional program and the differential transcriptional regulation of genes for BCL-2 family members depend on p53.

Based on the findings from the RNA-seq analysis (Figure 3C), we next investigated whether the observed transcriptional changes in the genes encoding the pro-survival proteins BCL-XL and MCL-1 were reflected at the protein level following p53 activation. Intracellular flow cytometric analysis revealed that treatment with nutlin-3a significantly increased BCL-XL protein levels in MDFs, BMDMs, and activated T as well as activated B cells, consistent with the transcriptional induction of their genes, whereas no such increase was observed in thymocytes or pre-B cells (Figure 3D; Supplementary Figure 6). MCL-1 protein levels were significantly elevated in BMDMs and activated T cells after treatment with nutlin-3a, but no significant upregulation in MCL-1 levels were seen in the other cell types examined (Figure 3E; Supplementary Figure 6).

### High basal pro-survival BCL-2 protein abundance and BCL-XL upregulation in cells that survive after p53 activation

To validate the differential regulation of BCL-XL (and MCL-1) at the global protein level, we performed quantitative mass spectrometry based proteomic analysis following 24 h of treatment with nutlin-3a in the presence of the pan-caspase inhibitor Q-VD-OPh to prevent protein degradation owing to apoptosis. Four primary cell types representing distinct cellular outcomes were examined (Figure 4A). In MDFs, BMDMs, and activated T cells, numerous proteins were differentially expressed following p53 activation, whereas fewer changes were detected in thymocytes (Figure 4B–4E).

**Figure 4.**
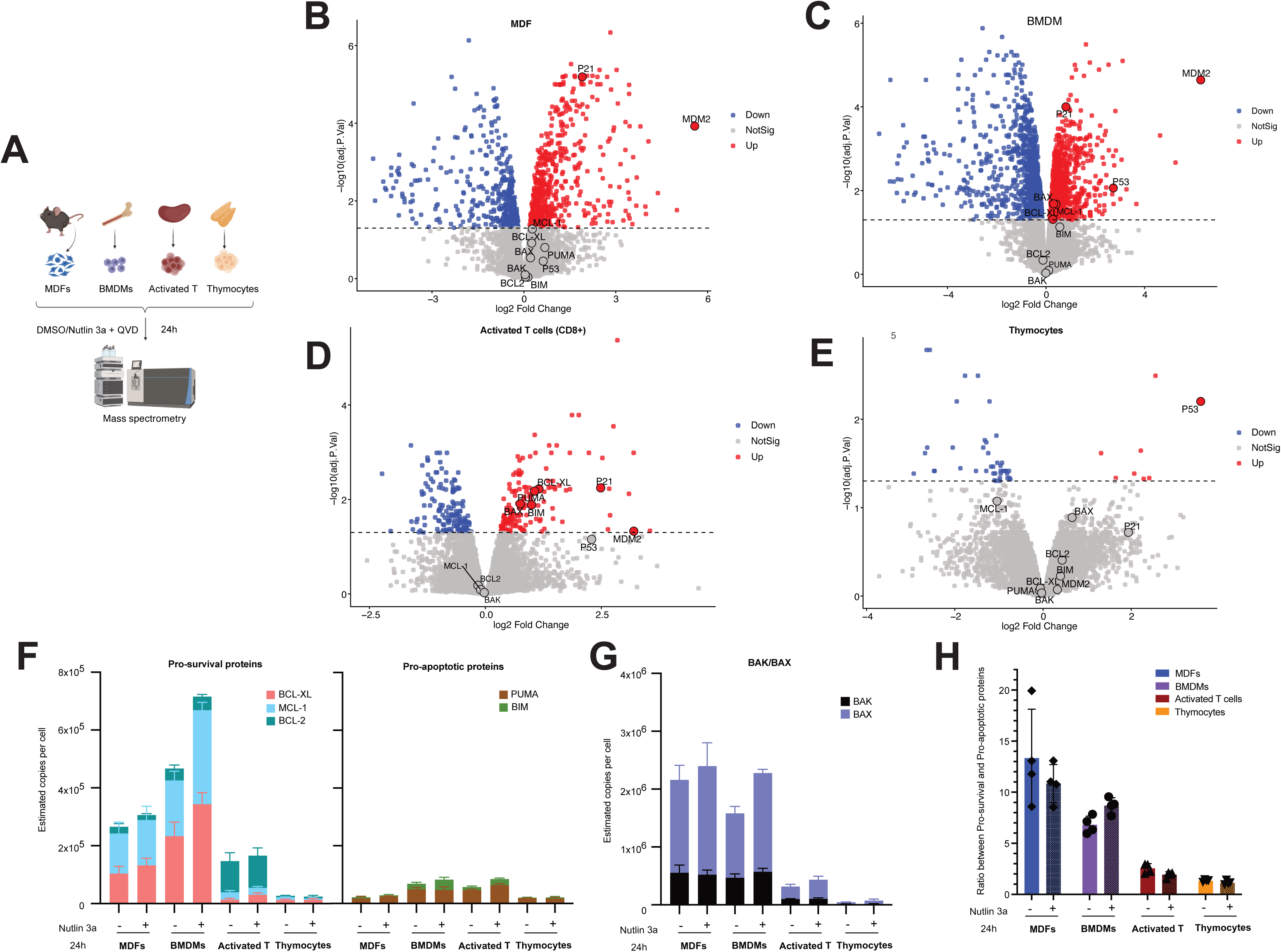
BCL-XL (*Bcl2l1*) is increased at the protein level in mass spectrometry analysis and high basal pro-survival BCL-2 protein levels are associated with cell survival after p53 activation. **A:** Schematic overview of the mass spectrometry experimental design. Proteins were isolated from MDFs, BMDMs, activated T cells, and thymocytes from wild-type mice following 24 h treatment with DMSO or 10 µM nutlin-3a in the presence of 25 µM QVD-OPH. **B–E:** Volcano plots showing changes in the levels of the indicated proteins upon p53 activation by nutlin-3a in MDFs (B), BMDMs (C), activated T cells (D), and thymocytes (E). Cells were treated for 24 h with 10 µM nutlin-3a in the presence of 25 µM QVD-OPH. Highlighted are proteins involved in p53 signalling (MDM2), apoptosis (BAK, BAX, BCL-2, MCL-1, BCL-XL, BIM, PUMA), cell cycle and senescence (p21) and p53 itself. NOXA was undetectable in the single-search data analysis. **F:** Copy numbers of pro-survival proteins (MCL-1, BCL-XL, and BCL-2) and pro-apoptotic proteins (BIM and PUMA) in MDFs, BMDMs, activated T cells, and thymocytes before and after 24 h treatment with nutlin-3a were quantified using the histone ruler method (copy number per million cells). **G:** Copy numbers of apoptotic effector proteins BAK and BAX per million cells in MDFs, BMDMs, activated T cells, and thymocytes before and after 24 h treatment with nutlin-3a were quantified using the histone ruler method (copy number per million cells). **H:** Ratio of pro-survival BCL-2 proteins (BCL-XL, MCL-1 and BCL-2) to pro-apoptotic BH3-only proteins (BIM and PUMA) in MDFs, BMDMs, activated T cells, and thymocytes before and 24 h after treatment with nutlin-3a.

Although thymocytes were co-treated with Q-VD-OPh and viable cells sorted prior to mass spectrometry, the limited proteomic changes may reflect the rapid apoptotic process in this cell type. Importantly, p53 protein itself was robustly increased in thymocytes (Figure 4E), confirming effective p53 activation. Consistent with the RNA-seq data, canonical p53 targets including MDM2 and p21 proteins were significantly increased at protein level in MDFs, BMDMs, which survive and undergo cell cycle arrest, as well as in activated T cells after treatment with nutlin-3a (Figure 4B–4D). In thymocytes, both proteins were also increased but this did not reach statistical significance, likely reflecting their rapid apoptosis and consequent reduced protein production (Figure 4E).

Notably, MCL-1 and BCL-XL proteins were increased in MDFs and BMDMs (Figure 4B and 4C) but not in thymocytes, which undergo rapid apoptosis following p53 activation (Figure 4E). While this increases did not reach statistical significance in MDFs by mass spectrometry, the trend was consistent with the findings from the intra-cellular flow cytometric analysis (Figure 3D,E). This may indicate that the increase in BCL-XL may be relatively modest in MDFs that are not readily resolved in bulk proteomic analysis but can be identified by flow cytometry. In activated T cells, BCL-XL levels were significantly increased upon p53 activation (Figure 4D), consistent with the FACS data (Figure 3E). Overall, increases in pro-survival protein BCL-XL were observed across cell types that exhibit survival or mixed outcomes following p53 activation but not in cells undergoing rapid apoptosis.

To exclude the possibility that the inability to see an increase in BCL-XL in thymocytes was merely due to their rapid apoptotic death, we generated haematopoietic chimeras by transplanting foetal liver cells, a rich source of haematopoietic stem and progenitor cells (HSPCs) from E14.5 *Bak^−/−^Bax^−/−^*embryos (C57BL/6-Ly5.2) into lethally C57BL/6-Ly5.1 recipient mice (Supplementary Figure 7A). Eight weeks post-transplantation, >95% donor cell contribution (Ly5.2+) to the haematopoietic compartment was confirmed by flow cytometry. Consistent with BAX and BAK having essential overlapping roles in the execution of apoptosis (Lindsten, Ross et al. 2000, Ke, Vanyai et al. 2018), *Bak^−/−^Bax^−/−^* thymocytes did not die after treatment with nutlin-3a (Supplementary Figure 7B). A subtle increase in BCL-XL was observed in *Bak^−/−^Bax^−/−^*thymocytes after treatment with nutlin-3a compared to wild-type thymocytes, and the reduction in MCL-1 was attenuated (Supplementary Fig. 7C–D). As these changes were not observed at the mRNA level (Figure 3C), they likely reflect post-transcriptional regulation and were associated with reduced apoptosis. BAX/BAK double-deficient activated T and B cells also exhibited markedly enhanced survival (∼90%) (Supplementary Figure 7E and 7I) but still underwent cell cycle arrest following p53 activation (Supplementary Figure 7F and 7J). Increases in MCL-1 (Supplementary Figure 5G and 5K) and BCL-XL (Supplementary Figure 5H and 5L) levels in these *Bax^-/-^Bak^-/-^*cells were similar to those observed in their wild-type counterparts after treatment with nutlin-3a. This supports the notion that BCL-XL upregulation after p53 activation is a predetermined response rather than a consequence of cell survival and proliferation arrest.

**Figure 5.**
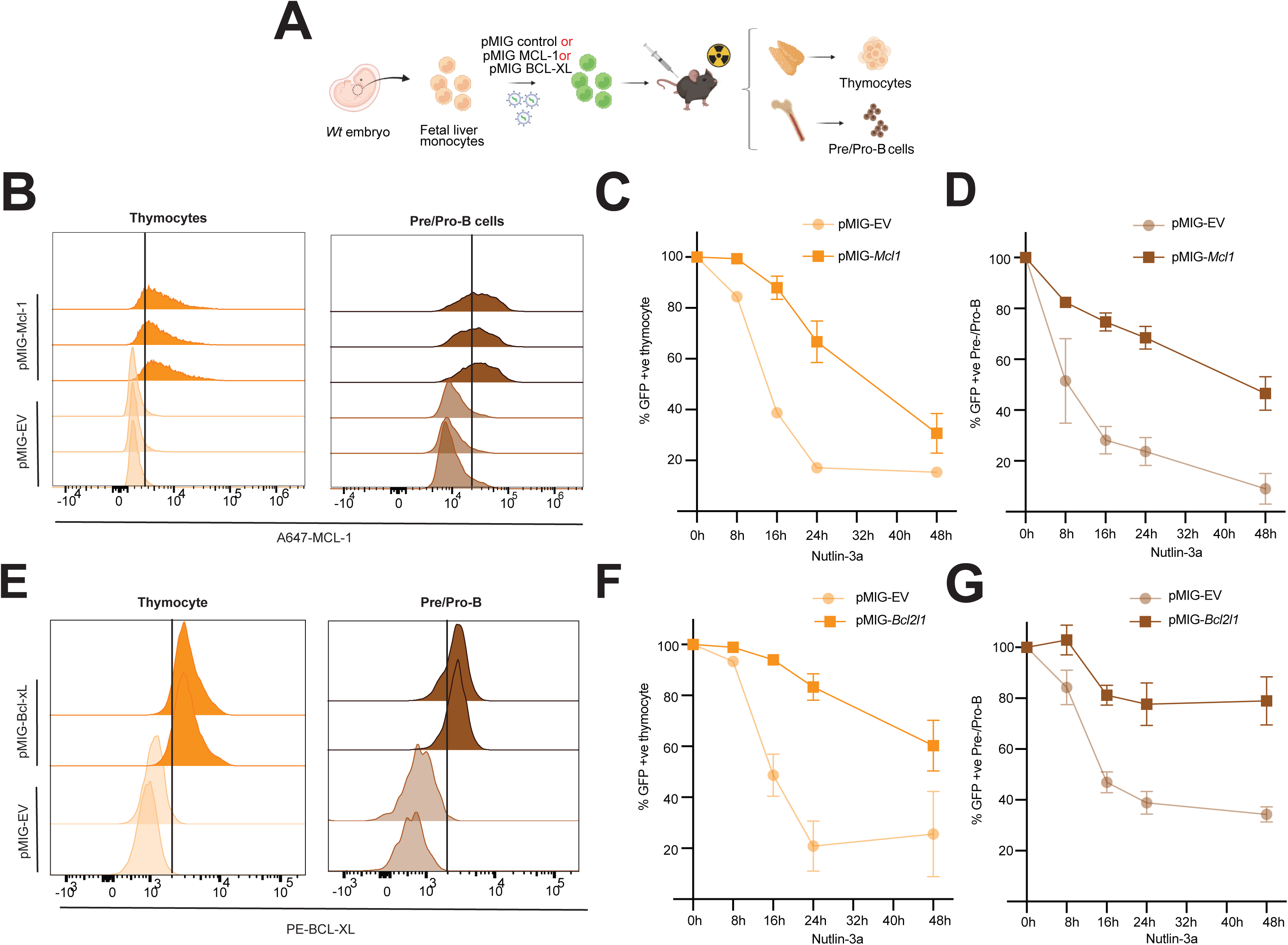
Overexpression of MCL-1 or BCL-XL confers resistance to p53-induced apoptosis in thymocytes and pre-B cells. **A:** Schematic illustrating the generation of thymocytes and pre-B cells overexpressing BCL-XL or MCL-1. Foetal liver cells from E14.5 C57BL/6-Ly5.2 embryos, a rich source of haematopoietic stem/progenitor cells (HSPCs) were retrovirally transduced with an MCL-1 overexpression construct (pMIG-*Mcl1*), a BCL-XL overexpression construct (pMIG-*Bcl2l1*) or an empty vector control (pMIG). Transduced cells were transplanted into lethally irradiated C57BL/6-Ly5.1 recipient mice to reconstitute their haematopoietic system. **B:** Representative flow cytometry histograms showing MCL-1 expression in thymocytes and pre-B cells isolated from mice reconstituted with a haematopoietic system transduced with a control virus (pMIG) or an MCL-1 overexpression construct (pMIG-*Mcl1*). Data are representative of n = 3 independent mice. **C** and **D:** Thymocytes (**C**) and pre-B cells (B220⁺IgM⁻) (**D**) overexpressing MCL-1 or an empty vector (EV) control were treated with 10 µM nutlin-3a for 8, 16, 24, and 48 h. Cell survival was measured as % GFP⁺ cells relative to DMSO controls. Data are presented as mean ± SD (n = 3 biological replicates). **E:** Representative flow cytometry histograms showing BCL-XL expression in thymocytes and pre-B cells harvested from mice reconstituted with a haematopoietic system transduced with a control virus (pMIG) or a BCL-XL overexpression construct (pMIG-*Bcl2l1*). Data are representative of n = 3 independent mice. **F** and **G:** Thymocytes (**F**) and pre-B cells (B220⁺IgM⁻) (**G**) overexpressing BCL-XL or an empty vector (EV) control were treated with 10 µM nutlin-3a for 8, 16, 24, and 48 h. Cell survival was measured as % GFP⁺ cells relative to DMSO controls. Data are presented as mean ± SD (n = 3 biological replicates).

The fate of cells after p53 activation, i.e. whether they will survive or undergo apoptosis, is likely determined not only by the increase in pro-survival BCL-2 proteins and pro-apoptotic BH3-only proteins but also by the starting (basal) levels of these proteins. Therefore, we measured the basal levels of pro-survival and pro-apoptotic members of the BCL-2 protein family across a range of cell types. Absolute protein copy numbers per cell were estimated using the proteomic histone ruler approach (Wiśniewski, Hein et al. 2014). Strikingly, basal levels of the pro-survival proteins BCL-XL, MCL-1 and BCL-2 were substantially higher in MDFs and BMDMs, which survive after p53 activation, compared with thymocytes, which die rapidly (Figure 4F). Activated T cells displayed intermediate levels, consistent with their mixed cellular outcomes. In contrast, basal levels of BH3-only proteins such as PUMA and BIM did not differ markedly across cell types with opposing fates (Figure 4F). Although BAK and BAX are present at high levels in all cell types examined (Figure 4G), these effectors require activation to execute apoptosis (Newton, Strasser et al. 2024). Therefore, the ratio of total pro-survival proteins to total pro-apoptotic BH3-only proteins was determined. This ratio was highest in MDFs and BMDMs, intermediate in activated T cells, and lowest in thymocytes. These findings indicate that intrinsic differences in basal pro-survival BCL-2 protein abundance and the balance between pro-survival BCL-2 proteins and pro-apoptotic BH3-only proteins may determine cell fate following p53 activation.

### BCL-XL and MCL-1 are critical for cell survival after p53 activation

Cell survival is safeguarded by BCL-XL, MCL-1 and other pro-survival BCL-2 proteins (Czabotar, Lessene et al. 2014, Newton, Strasser et al. 2024). Therefore, and based on the findings reported above, we hypothesised that if the low basal levels of pro-survival proteins, and their failure to increase BCL-XL and MCL-1 were the reasons why thymocytes and pre-B cells die after p53 activation, engineered expression of these pro-survival proteins would be expected to protect them from nutlin-3a induced killing. Indeed, when HSPCs from C57BL/6-Ly5.2 mice were retrovirally transduced with BCL-XL or MCL-1 expression constructs and then transplanted into lethally irradiated C57BL/6-Ly5.1 recipients (Figure 5A), the donor HSPC (i.e. Ly5.2+) derived thymocytes and pre-B cells showed increased expression of MCL-1 or BCL-XL as detected by intra-cellular FACS analysis (Figure 5B and 5E) and were much more resistant to nutlin-3a induced apoptosis compared to their control counterparts containing an empty vector (Figure 5C, D, F, G).

Conversely, we predicted that if the high basal protein levels of BCL-XL and MCL-1, together with the increase in BCL-XL (and MCL-1) are essential for the survival of MDFs and BMDMs after p53 activation, the genetic reduction or pharmacological inhibition of these pro-survival proteins should shift the cellular outcome from proliferation arrest with survival toward apoptotic cell death after treatment with nutlin-3a. Indeed, while treatment with nutlin-3a alone did not markedly reduce the viability of MDFs and BMDMs, their killing was markedly increased by adding the combination of intermediate doses of the BCL-XL inhibitor A1331852 (Lessene, Czabotar et al. 2013) and the MCL-1 inhibitor S63845 (Kotschy, Szlavik et al. 2016) (Figure 6A,C). Consistently, this combination of BH3 mimetics shifted the fate for MDFs and BMDMs from cell cycle arrest toward apoptosis, as evidenced by an increased sub-G1 population compared to treatment with nutlin-3a alone (Figure 6B, D and Supplementary Figure 7A, B). Activated T cells and B cells, which display mixed cellular outcomes following nutlin-3a treatment, were also sensitised to apoptosis by inhibition of BCL-XL or MCL-1, particularly when the two BH3 mimetics were combined. This resulted in increased cell death and an expanded sub-G1 population (Figure 6E-G and Supplementary Figure 7C, D). These findings support the notion that pre-exisiting (basal) levels and induction of BCL-XL and MCL-1 determine cell fate after p53 activation.

**Figure 6.**
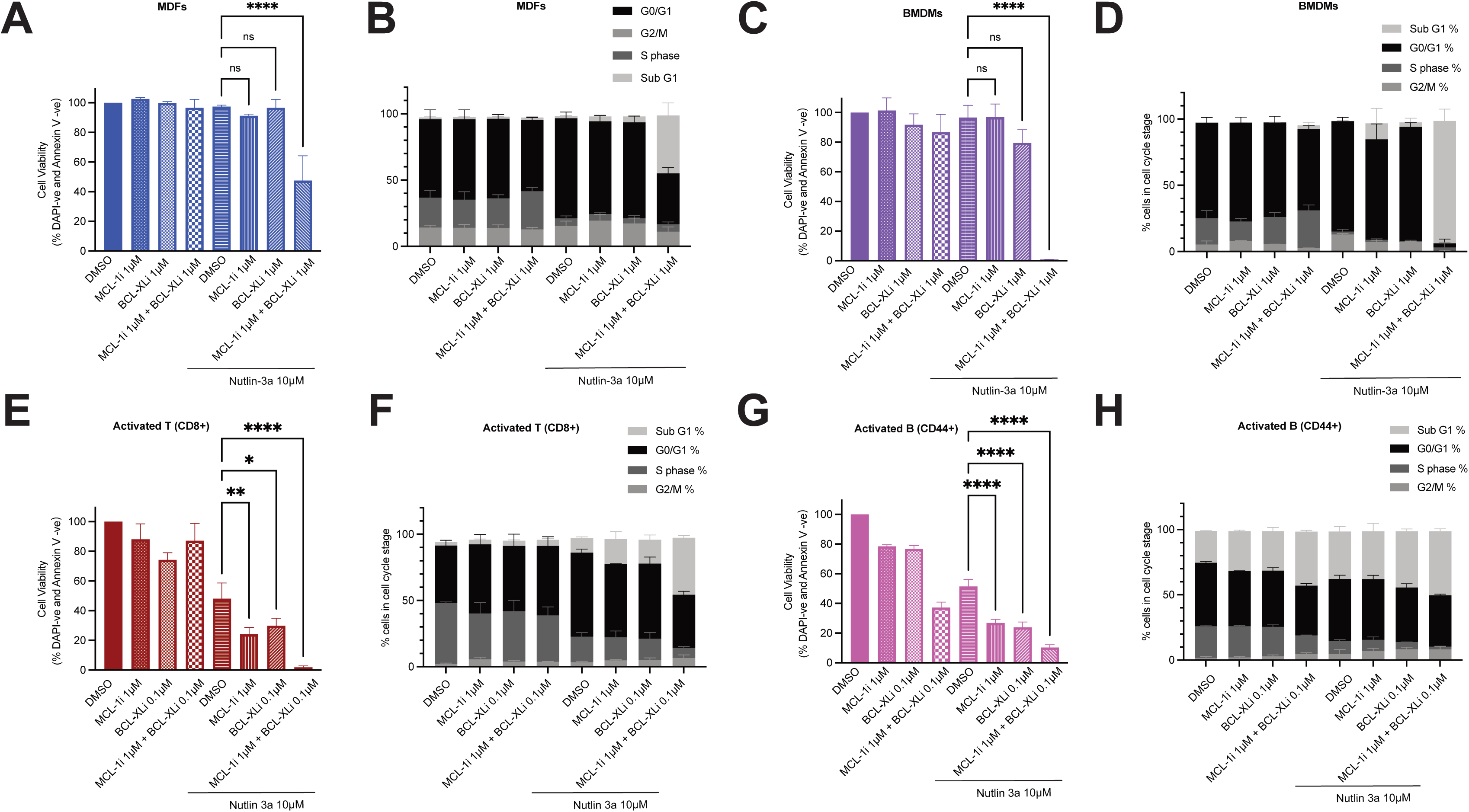
Pharmacological inhibition of BCL-XL and/or MCL-1 converts cell fate after p53 activation from survival with proliferation arrest to apoptotic death. **A-H:** MDFs (**A** and **B**), BMDMs (**C** and **D**), activated T cells (**E** and **F**) and activated B cells (**G** and **H**) were treated for 24 h with 1 µM MCL-1 inhibitor (S63845), 1 µM BCL-XL inhibitor (A-1331852), 10 µM nutlin-3a or the indicated combinations of agents with BrdU added to the cultures for the last 4h. Apoptosis (**A**, **C**, **E and G**) was assessed by staining with Annexin-V and DAPI followed by flow cytometry (Annexin V-DAPI-cells were deemed live cells). Cell cycle distribution (**B**, **D**, **F and H**) was determined by staining for BrdU incorporation and DNA content (7-AAD) and cells were divided into the following stages of the cell cycle: sub-G1 (apoptotic), G0/G1, S, or G2/M. Data are presented as mean ± SD (n = 3 biological replicates). Statistical significance was determined using two-way ANOVA with Dunnett’s multiple comparisons test (P < 0.05; P < 0.01; P < 0.001; P < 0.0001; ns, not significant).

### BCL-XL and MCL-1 protect human cancer cells from p53-induced apoptosis

Having established that elevated pro-survival BCL-XL (and MCL-1) protein levels with a high pro-survival BCL-2 protein to pro-apoptotic BH3-only proteins ratio determines cell fate after p53 activation in murine primary cells, we next examined whether this cell fate determining mechanism also operates in human cancer cells expressing wild-type p53. Upon treatment with nutlin-3a, the lung adenocarcinoma cell line A549, colon cancer cell line RKO and non-small cell lung cancer cell line H460 exhibited minimal apoptosis, whereas the human B cell lymphoma cell line DOHH2 showed extensive loss of viability (Figure 7A). The human osteosarcoma cell line SJSA-1 displayed an intermediate phenotype (Figure 7A). Consistent with our findings from primary murine cells, quantitative proteomic analysis revealed that the human cancer cell lines that did not die after p53 activation (A549, RKO and H460) exhibited a much higher basal ratio of pro-survival BCL-2 proteins to pro-apoptotic BH3-only proteins compared to apoptosis-sensitive lines (DOHH2) (Figure 7B). Notably, BCL-XL protein levels were significantly increased following treatment with nutlin-3a in A549, RKO and H460 cells. MCL-1 upregulation was observed in RKO and H460 cells but not in DOHH2 cells, based on both flow cytometry and mass spectrometry analysis (Figure 7C–G; Supplementary Figure 9A, B). Inhibition of BCL-XL and/or MCL-1 markedly enhanced nutlin-3a induced apoptosis in A549, RKO, H460 and SJSA1 cells (Figure 7H–K), converting a predominantly cell cycle arrest/cell survival outcome into robust apoptotic cell death. These findings demonstrate that high pre-exisitng (basal) levels of pro-survival BCL-2 proteins safeguard the survival of human cancer cells during p53 activation.

**Figure 7:**
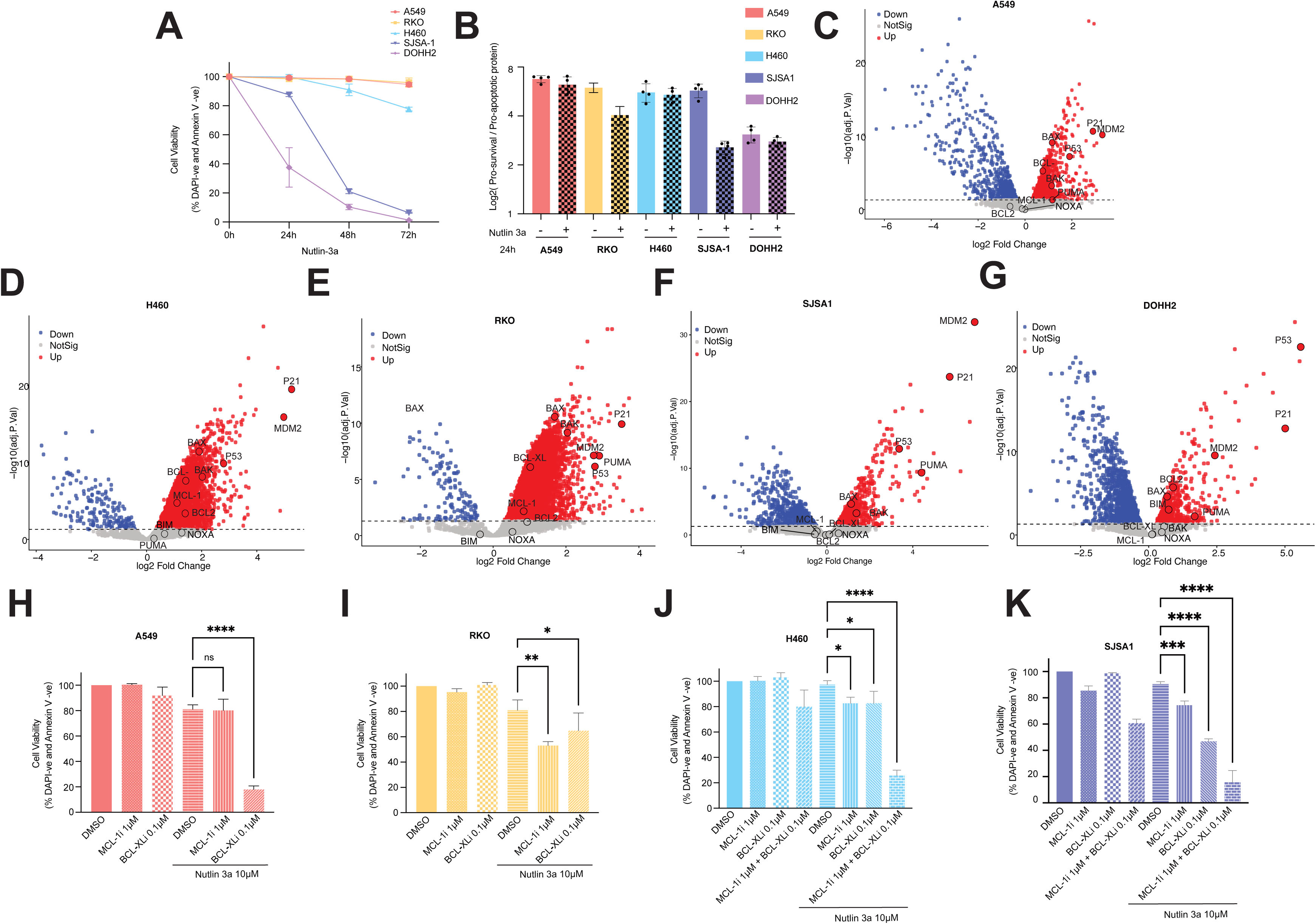
High ratio of pro-survival BCL-2 protein versus pro-apoptotic BH3-only proteins and BCL-XL upregulation in cancer cell lines resistant to p53 activation induced apoptosis. **A**: Cancer cell lines expressing wt p53 (A549, RKO, H460, SJSA-1 and DOHH2) were treated with 10 µM nutlin-3a for 24 h, 48 h and 72 h. Apoptosis was assessed by staining with Annexin-V and DAPI followed by flow cytometry (Annexin V-DAPI- cells were deemed live cells). Data are presented mean ± SD (n = 3 biological replicates). **B**: Ratio of pro-survival BCL-2 proteins (BCL-XL, MCL-1 and BCL-2) to pro-apoptotic BH-3 only proteins (BIM, PUMA and NOXA) in A549, RKO, H460, SJSA-1and DOHH2 before and after 24 h treatment with nutlin-3a. **C - G**: Volcano plots showing changes in protein levels upon p53 activation by nutlin-3a in A549 (**C**), H460 (**D**), RKO (**E**), SJSA1 (**F**) and DOHH2 (**G**). Cells were treated for 24 h with 10 µM nutlin-3a in the presence of 25 µM QVD-OPH. Highlighted are proteins involved in p53 signalling (MDM2), apoptosis (BAK, BAX, BCL-2, MCL-1, BCL-XL, BIM, PUMA and NOXA), cell cycle and senescence (p21) and p53 itself. **H-K:** Inhibition of BCL-XL and or MCL-1 enhances p53-induced apoptosis in A549 (**H**), RKO (**I**), H460 (**J**) and SJSA1 cells (**K**). Cells were treated for 24 h with 1 µM MCL-1 inhibitor (S63845), 1 µM BCL-XL inhibitor (A-1331852), 10 µM nutlin-3a, or the indicated combinations. Apoptosis was analysed as described in (**A**). Data are presented as mean ± SD (n = 3 biological replicates). Statistical significance was determined using two-way ANOVA with Dunnett’s multiple comparisons test (P < 0.05; P < 0.01; P < 0.001; P < 0.0001; ns, not significant).

### Predicting cell fate after p53 activation using the pro-survival BCL-2 to pro-apoptotic BH3-only protein ratio in human cancer cell lines

Our data show that a high basal pro-survival BCL-2 to pro-apoptotic BH3-only protein ratio safeguards the survival of both non-transformed murine cells and several human cancer cell lines during p53 activation. We wanted to explore whether this relationship could be extended to a broader range of human cancer cells and whether we could generate a model that predicts their fate after p53 activation. Only few proteomic data are available from human cancers. Therefore, we analysed transcriptomic data from the DepMap database.

For each cell line, mRNA expression levels of key pro-survival genes (*MCL1*, BCLXL/*BCL2L1* and *BCL2*) and critical pro-apoptotic BH3-only genes (*PUMA/BBC3*, *BIM* /*BCL2L11* and *NOXA/PMAIP1*) were obtained. Gene expression values were represented as log2(TPM+1). A composite expression score for each group was calculated by summing the log2(TPM+1) values of the corresponding genes. The pro-survival BCL-2 family gene to pro-apoptotic BH3-only gene ratio was then derived from these composite scores. Substantial variability in both pro-survival BCL-2 family gene and pro-apoptotic BH3-only gene expression was observed across cancer cell lines (Figure 8A). Consistent with our above reported findings (Figure 7B), cancer cell lines that survive and undergo cell cycle arrest after p53 activation (A549, RKO and H460) exhibited a higher basal pro-survival BCL-2 family gene to pro-apoptotic BH3-only gene ratio compared to the cancer cell line that undergoes apoptosis (DOHH2) (Figure 8B). This supports the concept that this ratio can predict cell fate after p53 activation. To predict the outcome of human cancer cells to p53 activation, we established a logistic regression model using experimentally measured apoptosis fractions (Figure 7A) and the corresponding pro-survival BCL-2 family member to pro-apoptotic BH3-only protein ratios calculated from log2(TPM) values (Figure 8C). The model generated a predicted probability of apoptosis (0–100%) for each cancer cell line based on its basal ratio. This model was subsequently applied to the broader panel of DepMap human cancer cell lines to estimate their likelihood of undergoing apoptosis upon p53 activation (Figure 8D, E). Consistent with the predictions from this model, cell lines with a high basal ratio (HEPG2 and MCF7) displayed a resistant phenotype, whereas those with relatively low ratios (OCI-AML2, OCI-AML3, MV4-11, SHSY5Y and MOLM13) underwent pronounced apoptosis following treatment with nutlin-3a (Figure 8F). These findings demonstrate that the model we developed can faithfully predict the outcome of human cancer cells to p53 activation.

**Figure 8:**
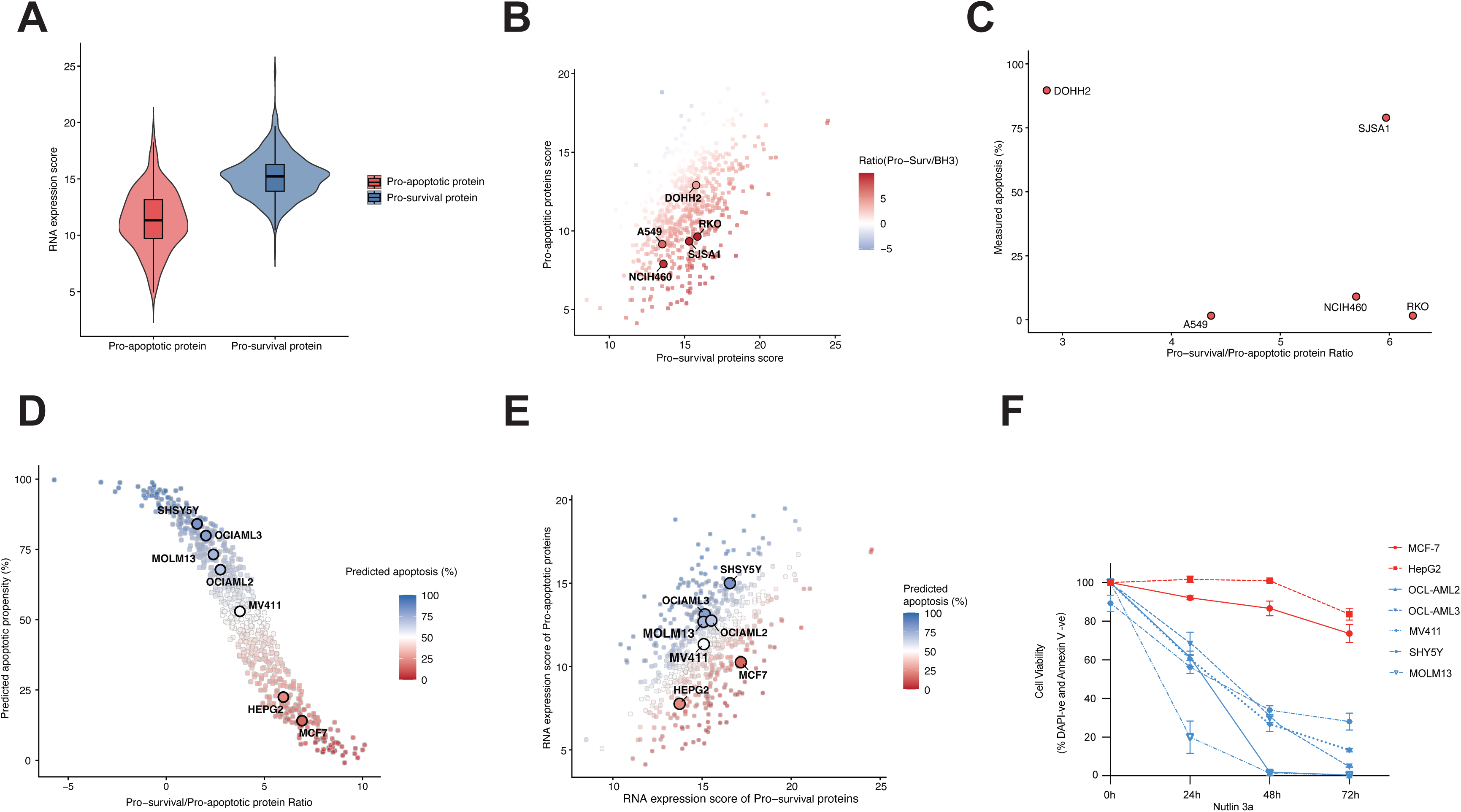
The ratio of pro-survival BCL-2 proteins to pro-apoptotic BH3-only proteins predicts propensity to undergo apoptosis after p53 activation across a broad range of human cancer cell lines. **A:** Violin and box plots showing the distribution of aggregated mRNA expression levels of pro-survival BCL-2 genes and pro-apoptotic BH3-only genes across human cancer cell lines harbouring wt p53from DepMap database. RNA expression scores were calculated by summing log_2_(TPM+1) values of pro-survival BCL-2 protein encoding genes (*BCL2L1/BCL-XL, MCL-1* and *BCL2*) and pro-apoptotic BH3-only genes (*PUMA/ BBC3*, *BIM/BCL2L11*and *NOXA/PMAIP1*). **B:** Distribution of wt p53 *human* cancer cell lines based on composite expression scores of pro-survival BCL-2 protein encoding genes and pro-apoptotic BH3-only protein encoding genes from the DepMap database. Each point represents a single human cancer cell line, and the colour indicates the pro-survival BCL-2 to pro-apoptotic BH3-only protein encoding mRNA ratio. Cell lines experimentally tested in Figure 7A are highlighted. **C:** Relationship between the basal pro-survival BCL-2 protein to-pro-apoptotic BH3-only protein expression ratio and experimentally measured apoptosis following p53 activation by 24 h treatment with nutlin-3a in the human cancer cell lines tested in Figure 7A. **D:** Predicted probability of undergoing apoptosis after p53 activation as a function of the pro-survival BCL-2 protein to-pro-apoptotic BH3-only protein ratio based on a logistic regression model generated from the experimentally measured data. Selected human cancer cell lines are highlighted for experimental validation. **E:** Predicted probability of undergoing apoptosis after p53 activation across wt p53 human cancer cell lines mapped onto the pro-survival BCL-2 protein versus pro-apoptotic BH3-only protein expression landscape. The colour scale represents predicted probability of cells undergoing apoptosis after p53 activation. Selected cancer cell lines are highlighted for experimental validation. **F:** Experimental validation of predicted apoptotic responses following p53 activation in the indicated human cancer cell lines from panel D and E. The human cancer cell lines (MCF-7, HepG2, OCL-AML2, OCL-AML3, MV411, SHY5Y and MOLM13) were treated with 10 µM nutlin-3a for 24 h, 48 h and 72 h. Apoptosis was assessed by staining with Annexin-V and DAPI followed by flow cytometry (Annexin V-DAPI- cells were deemed live cells). Data are presented as mean ± SD (n = 3 biological replicates).

## Discussion

In this study, we performed mutli-omics analyses across a range of primary murine cell types (MDFs, BMDMs, activated T, activated B, thymocytes and pre-B) and human cancer cell lines with opposing cellular outcomes upon p53 activation. Our findings reveal that how cells determine cell fate upon p53 activation is not solely dependent on selective induction of p53 target genes encoding mediators of cell cycle arrest or initiators of apoptosis, respectively. Consistent with a few published studies (Kenzelmann Broz, Spano Mello et al. 2013, Valente, Aubrey et al. 2016, Moyer, Wasylishen et al. 2020), our data indicate that the induction of cell cycle arrest inducing gene *P21*(*Cdkn1a)* and the critical pro-apoptotic gene *Puma(Bbc3)* after p53 activation are not mutually exclusive; both genes were induced regardless of whether cells underwent proliferation arrest/senescence or apoptosis. Genetic deletion of *P21*(*Cdkn1a)* did not substantially alter the overall cellular outcomes induced by p53 activation, suggesting that p21-mediated cell cycle arrest alone is insufficient to explain the different cellular outcomes induced by p53 activation between different cell types.

Instead, our data reveal that cell fate following p53 activation is determined by the pre-existing (basal) levels of pro-survival BCL-2 proteins. Cells undergo cell cycle arrest/senescence following p53 activation exhibited high basal expression of pro-survival BCL-2 proteins, resulting in a higher pro-survival BCL-2 protein versus pro-apoptotic BH3-only protein ratio. In contrast, cells with relatively low pro-survival BCL-2 protein levels and a low ratio undergo apoptosis after p53 activation. These findings support a model in which the balance of BCL-2 family proteins establishes the apoptotic threshold that determines whether cells undergo reversible cell cycle arrest or commit irreversibly to apoptotic death after p53 activation. In addition to differences in the basal levels of pro-survival BCL-2 proteins, we found that p53 activation can upregulate pro-survival BCL-XL (*Bcl2l1*) at both the transcript and protein level in cells that survive after p53 activation., Genome-wide CRISPR/Cas9 screening identified that this increase in BCL-XL is p53-dependent (Supplementary figure 10). MCL-1 was similarly upregulated in several cell types that survived after p53 activation. These observations suggest that, in addition to inducing canonical pro-apoptotic target genes such as *Bbc3/Puma*, *Pmaip1/Noxa*, and *Bax*, p53 activation can also promote pro-survival responses in certain cellular contexts. Of note, previous work (Kenzelmann Broz, Spano Mello et al. 2013) and our cut & run experiments showed that p53 can bind to target sites in the *Bcl2l1* gene encoding BCL-XL (Supplementary figure 11). Combined with high basal pro-survival protein level, the upregulation of pro-survival proteins BCL-XL and MCL-1 may further raise the apoptotic threshold and prevent cells from apoptosis following p53 activation.

Our findings also have potential therapeutic implications. Pharmacological inhibition of BCL-XL and/or MCL-1 converted p53-induced cell cycle arrest into apoptosis across multiple primary murine cell types and also in certain human cancer cell lines expressing wild-type p53. Consistent with our findings, combination strategies involving p53 activation alongside inhibition of pro-survival BCL-2 family proteins have demonstrated promising anti-tumour activity in several cancer models harbouring wild-type p53 (Bharti, Watkins et al. 2022, Bell, Blair et al. 2024). However, the mechanistic basis underlying these responses was not fully explained. For example, it was reported that combined MDM2 or AURKA inhibitor together with pro-survival protein inhibition preferentially promotes apoptosis over cell cycle arrest in melanoma and acute lymphoblastic leukemia model and was associated with reduced levels of p21 protein (Bharti, Watkins et al. 2022, Bell, Blair et al. 2024). However, *P21*(*CDKN1A)* transcription was not significantly different compared with p53 activation agent treatment on its own. This may indicate that reduced p21 protein level may be a consequence rather than a driver of apoptosis. Since p21 is a relatively short-lived protein (Sheaff, Singer et al. 2000), it is likely rapidly degraded during apoptotic cell death. Our findings support this interpretation as MDF and BMDM from *P21*(*Cdkn1a)^-/-^* mice did not shift the cell fate from proliferation arrest toward apoptosis upon p53 activation. Regardless, collectively, these findings provide a mechanistic rationale for combining p53 activation therapies with inhibition of pro-survival proteins to effectively killing cancer cells with wild-type p53.

Our findings indicate that the ratio between pro-survival BCL-2 protein *versus* pro-apoptotic BH3-only proteins can be inferred from RNAseq data and used to provide a predictive framework of cellular outcomes following p53 activation. Although predictions of propensity of cancer cells to undergo apoptosis can also be done by BH3 profiling (Chonghaile, Sarosiek et al. 2011, Montero, Sarosiek et al. 2015), this requires access to live tumour cells from patients and extensive *in vitro* experimentation. In contrast, our model to predict cell fate only needs RNAseq data which are becoming increasingly available in a real life clinical setting. It must however be noted that our predictive model was derived from cancer cell line RNAseq datasets and *in vitro* tests. Future studies will be required to determine how well our model will predict *in vivo* tumour responses and in human patients.

In conclusion, our studies using integrated multi-omics analyses across a range of primary murine cell types and human cancer cell lines, reveal that the basal levels of pro-survival BCL-2 proteins as well as their further transcriptional induction and their ratio to pro-apoptotic BH-3-only proteins establishes the apoptotic threshold that determine whether cells survive and undergo proliferation arrest/senescence or undergo apoptotic death. These findings suggest p53 creates a stress response state, where the pre-existing state of the BCL-2 family network determines whether cells cross the apoptotic threshold. This framework provides mechanistic insight into the longstanding problem of p53 cell fate determination and suggests a potential strategy for predicting and therapeutically modulating responses to p53 activation in cancer.

## Method

### Mouse strains

All animal experiments were conducted with the approval of the WEHI Animal Ethics Committee following the guidelines set out by the Melbourne Directorate Animal Ethics Committee. *Trp53^−/−^* mice have been previously described (Jacks, Remington et al. 1994). They were originally generated on a mixed C57BL/6x129SV background but had been backcrossed with C57BL/6 mice for >20 generations prior to the commencement of the studies described here.

*p21^-/-^* mice (Deng, Zhang et al. 1995), *Bak^-/-^Bax^+/-^* (Lindsten, Ross et al. 2000) and Cas9*^KI/KI^* mice (Chu, Weber et al. 2016) were generated on a C57BL/6 background and have been previously described. *Puma-tdTomato*;*p21-IRES-GFP* double reporter mice were developed by intercrossing *p21-IRES-GFP* and *Puma-tdTomato* reporter strains which have been previously described (Lieschke, Thomas et al. 2024).

### Genotyping of mice by PCR

Crude DNA was extracted from either a tailpiece or an ear clip obtained at weaning, using the DirectPCR Lysis buffer (Viagen #102-T) with 0.32 U/mL proteinase K (Sigma-Aldrich #P4850) at 56 °C for a minimum of 4 h until completion of tissue digestion. Proteinase K was inactivated at 86 °C for 20 min. PCR was performed using GoTaq Green Master Mix (Promega #M7123) followed by the protocol descripted previously (Lindsten, Ross et al. 2000, Lieschke, Thomas et al. 2024). PCR products were run on a 2% agarose gel in 1X TAE buffer (Sigma-Aldrich #T9650 diluted with water) containing 0.5 µg/mL ethidium bromide at 100 V for 40 min and visualised using the ChemiDoc XRS+ in the ImageLab programme.

### Mouse dermal fibroblasts (MDFs) generation

MDFs were generated from the tails of 8-12-week-old mice. Mice were euthanised by cervical dislocation by an experienced animal technician, or by CO2 inhalation. Whole tails were removed using surgical scissors and were placed in ice cold DMEM (Gibco, #11885-084). In the sterile hood, each tail was sprayed with 80% ethanol and then placed into a Petri dish (Corning, #430167). Bone was removed from the skin using tweezers and a surgical blade and the skin was digested in 2.1 U/mL dispase II (Sigma-Aldrich #D4693) in DMEM at 4°C overnight on a roller. The dermis was peeled from the epidermis, cut into smaller pieces using surgical scissors, and then placed into 0.42 mg/mL Collagenase IV (Sigma-Aldrich #C5138) in DMEM at 37°C for 1 h. The dermis was then gently passed through a 100 μm mesh cell strainer (Falcon, #352360) and collagenase was rinsed off using DMEM. Cells were plated in 2 wells of a 6-well plate in DMEM, supplemented with 100 U/mL penicillin, 100 mg/mL streptomycin (Sigma #p0781) and 10% heat-inactivated (HI) foetal calf serum (FCS). Cells were grown at 37°C in a humidified atmosphere containing 2% O_2_ 10% CO_2_ and were passaged every 2-3 days until natural cell senescence occurred (usually ∼2 weeks after original plating). All assays were performed on early passage (p3-5 MDFs).

### Bone marrow-derived macrophages (BMDMs) generation

A single cell suspension of bone marrow cells was generated by collecting and flushing two femora and two tibiae with Dulbecco’s Phosphate Buffered Saline (DPBS, Gibco #14190). The cell suspension was centrifuged at 1500 rpm/300 g for 5 min, followed by aspiration of the supernatant. Cells were resuspended in 8 mL DMEM (Gibco, #11885-084), supplemented with 100 U/mL penicillin, 100 mg/mL streptomycin (Sigma-Aldrich #P0781), 10% HI-FCS and 15% L929 supernatant (a source of macrophage colony-stimulating factor (MCSF); made in house, WEHI). 1 mL of cell suspension in addition to 7 mL fresh media was then plated across 8 non-coated 10 cm culture dishes (Corning, #430167). Cells were incubated at 37°C in a humidified atmosphere containing 10% CO₂ for 3 days undisturbed. After 3 days, 4 mL medium containing L929 cell supernatant was added and the cells were then left for another 4 days undisturbed before further experiments.

### Isolation and stimulation of primary T lymphoid cells

Murine spleens were gently passed through a 100 mm mesh cell strainer (Falcon, #352360) to generate single cell suspensions in DPBS (Gibco #14190) with 3% HI-FCS. Red blood cells were depleted by resuspending cells in Red Cell Lysis Buffer (made in house, WEHI) for 5 min at room temperature. The cells were washed in DPBS (Gibco #14190) with 3% HI-FCS and centrifuged at 1500 rpm/300 g for 5 min. Cells were then resuspended in RPMI 1640 medium supplemented with 100 U/mL penicillin, 100 mg/mL streptomycin (made in house, WEHI), 1x GlutaMAX (Gibco #35050-061), 1x sodium pyruvate (Gibco, #11360-070), 1x MEM non-essential amino acids (Thermo Fisher Scientific, #11140050), 1x HEPES (Gibco, #15630-080), 10% HI-FCS and 50 μM β-mercaptoethanol (BDH chemicals, Lot ZA3568599) in a 6-well plate (Corning Costar, #3516). Cells were then stimulated for 72 h with aCD3 (500 ng/mL, Clone 145-2C11, made in house, WEHI), aCD28 (500 ng/mL, Clone 37N51, made in house, WEHI) and mIL2 (1:100, made in house, WEHI), at 37°C in a humidified atmosphere containing 5% CO₂. After stimulation, cells were transferred to 10 mL tubes and centrifuged at 1500 rpm for 5 min. Supernatant was aspirated and cells were resuspended in fresh T cell medium, supplemented with fresh mIL2. Cells were then plated out for further experiments.

### Isolation and stimulation of primary B lymphoid cells

Murine spleens were gently passed through a 100 mm mesh cell strainer (Falcon, #352360) to generate single cell suspensions in DPBS (Gibco #14190) with 3% HI-FCS. Red blood cells were depleted by resuspending cells in Red Cell Lysis Buffer (made in house, WEHI) for 5 min at room temperature. The cells were washed in DPBS (Gibco #14190) with 3% HI-FCS and centrifuged at 1500 rpm/300 g for 5 min. Cells were then resuspended and stimulated with IL-4 (100 U/mL, made in house, WEHI), 10 μg/mL anti-CD40 antibody (FGK45, made in house, WEHI) and 20 μg/mL anti-IgM Fab2 antibody fragments (Jackson AffiniPure^TM^, #115-006-075) in RPMI 1640 medium supplemented with 100 U/mL penicillin, 100 mg/mL streptomycin (made in house, WEHI) in a 6-well plate (Corning Costar, #3516). Cells were then left undisturbed for 48 h at 37°C in a humidified atmosphere containing 5% CO₂ before further experiments.

### Thymocytes generation

Murine primary thymi were harvested and placed in 2mL DBPS (Gibco #14190) with 3% HI-FCS. Thymi were gently passed through a 100 μm mesh cell strainer (Falcon, #352360) to generate a single cell suspension of thymocytes in DPBS (Gibco #14190) with 3% HI-FCS. Red blood cells were depleted by resuspending cells in Red Cell Lysis Buffer (made in house, WEHI) for 5 min at room temperature. The cells were washed in DPBS (Gibco #14190) with 3% HI-FCS and centrifuged at 1500 rpm/300 g for 5 min. Cells were then resuspended in FMA medium consisting of high glucose DMEM supplemented with 10% HI-FCS, 100 μM L-asparagine (Sigma #A4284), 50 μM β-mercaptoethanol (Sigma #M3148), 100 U/mL penicillin and 100 mg/mL streptomycin (Gibco #15140122) for further experiments.

### Pre-B cells generation

Both femora and both tibias were collected, and bone marrow was flushed into DPBS (Gibco #14190) with 3% HI-FCS to generate a single cell suspension. Whole bone marrow was stained with biotinylated antibodies against Ter119 (WEHI), Mac1 (clone MI/70, WEHI) and Gr1 (clone RPB6-8C5, WEHI), before incubation with MagniSort Streptavidin Negative Selection Beads (Invitrogen #MSNB-6002), using a magnet to deplete the bone marrow of myeloid and erythroid cells. The remaining cells were stained with antibodies against B220 (clone RA3-6B2 Catalogue# 563708, BD Biosciences) and IgM (clone 5.1, WEHI) for 20 min on ice before being washed and resuspended in DPBS (Gibco #14190) with 3% HI-FCS and 1ug/mL propidium iodide (PI). Single B220+ IgM- PI- pre-B cells were sorted on an Aria Fusion cell sorter (Becton Dickinson). Collected pre-B cells were cultured in MEM-alpha medium (Gibco #32561-037) supplemented with 10 nM HEPES (Gibco#15630-80), 1 mM sodium pyruvate (Gibco # 11360-070), 50 uM beta-mercaptoethanol (Sigma-Aldrich # M-6250), 100U/mL penicillin, 100mg/mL streptomycin (Gibco # 15140-122) and 20% HI-FCS + 1:100 IL-7 supernatant (made in house) for further experiments.

### Human cancer cell lines and cell culture conditions

Human colorectal carcinoma cell line RKO (a kind gift from Prof. D. Huang, WEHI), lung adenocarcinoma cell line A549 (a kind gift from Prof. D. Huang, WEHI), hepatoblastoma cell line HepG2 (a kind gift from A/Prof. N.Y. Fu, WEHI) and neuroblastoma cell line SH-SY5Y (a kind gift from Prof. G. Dewson, WEHI) were cultured in DMEM supplemented with 10% HI-FCS. Non-small cell lung cancer cell line H460 (a kind gift from Prof. D. Huang, WEHI), breast cancer cell line MCF-7 (a kind gift from Prof. J. Visvader, WEHI), B-cell non-Hodgkin lymphoma cell line DoHH2 (a kind gift from Prof. D. Huang, WEHI), osteosarcoma cell line SJSA-1(a kind gift from Prof. D. Huang, WEHI) and AML cell lines (MOLM13, MV4;11, OCI-AML2 and OCI-AML3; a kind gift from Prof. A. Wei, WEHI) were cultured in RPMI-1640 supplemented with 10% HI-FCS. All cell lines were maintained at 37°C in a humidified atmosphere containing 5% CO₂ and were routinely tested for mycoplasma contamination using the Lonza MycoAlert™ Mycoplasma Detection Kit (LT07-318) according to the manufacturer’s instructions.

### Cell death assays

Murine cell types and human cancer cell lines were seeded into 24-well flat-bottom plates at 1 x 10^5^ cells/well and cultured with DMSO, Nutlin-3a (10 µM; MedChemExpress, #HY-10029), A-1331852 (0.1 µM; WEHI Chemical Biology Division), S-63845 (1 µM; Cat#A-6044, Active Biochem) or the indicated drug combination for durations specified in the corresponding figure legends. Following treatment, both viable and dead cells were harvested for analysis. Activated T cells and B cells were first stained with cell surface marker FITC-conjugated anti-CD8 antibody (1:400; WEHI) for activated T cells and Brilliant Violet (BV) 605-conjugated anti-B220 (1:200; BioLegend, #103244) for activated B cells on ice before viability assessment. Cells were then resuspended in 1× Annexin V binding buffer (prepared by diluting a 10× stock containing 25 mM CaCl₂, 1.4 M NaCl, and 0.1 M HEPES, pH 7.4, in distilled water) containing Annexin-647 (made in house, 1:4000), and DAPI (0.5 µg/mL). Cell viability (Annexin V/DAPI double-negative) were quantified on a LSR II or Fortessa X-20 flow cytometer (BD Biosciences). Data analysis was performed using FlowJo v10 (BD Biosciences) and Prism v9.

### Cell cycle arrest assays

5 x 10^5 MDFs and BMDMs were seeded per well in 6-well plates and allowed to adhere overnight at 37°C in a humidified atmosphere containing 10% CO₂. Next day, culture medium was replaced with fresh medium containing either DMSO (Sigma, D4540) or Nutlin-3a (10 μM; MedChemExpress, HY-10029). Activated T and B cells (5 x 10^5 per well) were seeded immediately in 6-well plates before treatment of DMSO (Sigma, D4540) or Nutlin-3a (10 μM; MedChemExpress, HY-10029). Cells were incubated for 24 h or 48 h, with BrdU (3 μg/mL; APC BrdU Flow Kit, BD Pharmingen™, #557892) added during the final 4 h of culture. Followed by the incubation, MDFs and BMDMs were dissociated with trypsin (Sigma, #T4174), combining the supernatant with the trypsinised fraction for analysis. Activated T cells and activated B cells were harvested directly and transferred to FACS tubes and then stained with cell surface marker FITC-conjugated anti-CD8 antibody (1:400; WEHI) for activated T cells and Brilliant Violet (BV) 605-conjugated anti-B220 (1:200; BioLegend, #103244) for activated B cells on ice before being washed and resuspended in DPBS containing 3% FCS. Cells were fixed and stained using the APC BrdU Flow kit (BD Pharmingen^TM^, #557892) according to the manufacturer’s instructions. Following staining, cells were resuspended in BD Perm/wash buffer with 7-AAD (2.5 µg/mL; supplied in APC BrdU Flow kit, #557892). Samples were analysed within 24 h on the LSR IIW flow cytometer (Becton Dickinson) at a flow rate of fewer than 400 events per second.

### qRT-PCR analysis

1 x 10^6 cells (MDFs, activated T cells and thymocytes) were treated with either DMSO or nutlin- 3a (10 μM; MedChemExpress, HY-10029) for the indicated times (2, 4, 6, 8, or 24 h). For activated T cells and thymocytes, pan-caspase inhibitor Q-VD-OPh (25 μM; MedChemExpress, HY-12305) was added to block cell death during treatment. Following the treatment, cell pellets were frozen in TRIzol (Thermo Fisher Scientific #15596018), and RNA extracted from these pellets, as per manufacturer’s instructions. Synthesis of cDNA was performed using the SuperScript III First Strand Synthesis system (Thermo Fisher Scientific #11904018 or #11752050), as per manufacturer’s instructions. qRT-PCR assays were performed using TaqMan Fast Advanced Master Mix (Thermo Fisher Scientific #4444557), and TaqMan Assay probes targeting *Hmbs* (Mm01143545_m1), *Bbc3* (Mm00519268_m1), and *Cdkn1a* (Mm00432448_m1) (all from Thermo Fisher Scientific), each according to the manufacturer’s instructions. All qRT-PCRs used 3 technical replicates per sample, and were performed across 3 biological replicates. All qRT-PCRs were run on a QuantStudio 12K Flex Real-Time PCR System (Thermo Fisher Scientific). Data were analysed using the ΔΔCt method, with each sample normalised to expression of the chosen housekeeping control gene (*Hmbs*) and then normalised to gene expression in their respective 0 hr samples.

### Flow cytometry for activation of reporter genes

15,000 adherent cells (MDFs and BMDMs) were plated in 12-well plates in their respective culture media and were left to adhere for overnight in an incubator with 10% CO_2_ at 37°C. Medium was then replaced with medium containing DMSO, or 10 10 μM Nutlin-3a. Cells were incubated for up to 72 h. 40,000 mitogen-activated B and T lymphoid cells were plated out in triplicate in 96-well flat-bottomed plates (Falcon, #353072) and incubated with the respective culture media containing either DMSO or 10 μM Nutlin-3a for up to 24h at 37°C and 5% CO_2_. 25 μM pan-caspase inhibitor Q-VD-OPh was added to block cell death during treatment. At the time of analysis, MDFs and BMDM were dissociated with trypsin. Both the trypsinised fractions and supernatant were kept for analysis. Cells were centrifuged at 1500 rpm/300 g for 5 min to remove medium and were resuspended in DPBS containing DAPI (0.5 µg/mL) to stain for dead cells. Flow cytometry for adherent cells was carried out every 24 h up to 72 h on an Analyser BD FACSymphony A3 (E4.51) to detect live/dead cells, and cells expressing the *p21*-IRES-GFP and *Puma*-tdTomato reporters. Activated T cells and activated B cells were transferred to 96-well round-bottomed plates (Falcon, #353077) and were centrifuged at 1500 rpm/300 g for 5 min. The supernatant was flicked off and the cells were stained with cell surface marker APC-conjugated anti-Thy1 antibody (1:50; WEHI) for activated T cells and A700-conjugated anti-B220 (1:400; BioLegend, #103232) for activated B cells on ice for 20 min before being washed and resuspended in DPBS containing DAPI (0.5 µg/mL). Flow cytometry was carried out on a Novocyte Penteon Analyser to detect live/dead cells, and cells expressing the *p21*-IRES-GFP and *Puma*-tdTomato reporters. Data analysis was performed using FlowJo v10 and Prism v9.

### RNA isolation and Illumina RNA sequencing library preparation

MDFs and BMDMs were plated at the desired cell density (1x10^5^ per well in 12-well plates for MDFs and 6x10^5^ per well in 6-well plates for BMDMs) and left for overnight to adhere. Pre-B cells (1x10^6^ cells per well), thymocytes (6x10^6^ cells per well), activated T cells (1x10^6^ cells per well) and activated B cells (1x10^6^ cells per well) were plated in 12-well plates. All the cells were treated with either DMSO or 10 μM nutlin-3a for 6h and incubated in the designated incubator. Q-VD-OPh (25 μM) was added to block cell death during treatment for the suspension cells. Following the treatment, cells were harvested in TRIzol lysis reagent (Invitrogen #15596026) and then stored at -80°C until RNA was extracted. For RNA sequencing, RNA was extracted using the Direct-zol RNA MicroPrep kit (Zymo #R2060), according to the manufacturer’s instructions, including an on-column DNAse digestion. Samples were run on an Agilent 2200 TapeStation using RNA Screen Tape (#5067-5576) to check the purity of the samples. RNA quality was deemed acceptable if two clear peaks were present, corresponding to the 18S and 28S ribosomal peaks. RNA sequencing (RNAseq) libraries were prepared using the TruSeq RNA Library Prep Kit (Illumina #RS-122-2001) as per the manufacturer’s instructions. 10-100 ng of total RNA was used as input to make the library. mRNA was isolated using polyA beads and was then converted into cDNA. Illumina adapters were ligated, and libraries were enriched using 15 PCR cycles of the PCR protocol prior to being run on a D1000 TapeStation tape to check library quality. Samples were pooled and run on a Next-Seq using paired-end chemistry, by Dr Stephen Wilcox in the WEHI genomics laboratory.

### RNA-seq data analysis

Samples originated from six cell types (MDFs, BMDMs, activated T cells, activated B cells, thymocytes, and pre-B cells), with three biological replicates per combination of cell type, genotype (WT and p53 KO), and treatment (Nutlin and DMSO), yielding 72 biological samples in total. Technical replicates from nine individual MDF, pre-B, and thymocyte samples were generated to increase the sequencing depth of the original libraries. Samples from BMDM and activated T and B cells were sequenced in separate batches with one cell type per batch. Samples from MDF, thymocyte, and pre-B cells were sequenced across three batches. Raw paired-end FASTQ files were aligned to the mouse reference genome mm39 using the align function from the Rsubread package (v2.24.0)(Liao, Smyth et al. 2019). All BAM files were coordinate-sorted during alignment by setting sortReadsByCoordinates = TRUE. Read counts were obtained for each gene using the featureCounts function from the Rsubread package, with a custom RefSeq strict annotation containing only exons from “BestRefSeq” or “RefSeq” or “Curated Genome” annotation sources (file 230411-mm39_RefSeqStrict_MaskY.saf.gz downloaded from URL https://bioinf.wehi.edu.au/Rsubread/annot/current).

Differential expression analyses of the gene read counts were conducted using the edgeR (v4.8.0)(Chen, Chen et al. 2025) and limma (v3.66.0)(Ritchie, Phipson et al. 2015) packages. First, read counts were organised into a single DGEList object using the featureCounts2DGEList function from the edgeR package, with samples from all six cell types combined. Technical replicates were summed prior to downstream analysis using edgeR’s sumTechReps function. Gene annotation was obtained from the NCBI database (file Mus_musculus.gene_info.gz downloaded from URL https://ftp.ncbi.nlm.nih.gov/gene/DATA/GENE_INFO/Mammalia on 3 February 2026), from which gene symbols and gene biotypes were extracted. Genes were filtered to retain only those located on autosomal or sex chromosomes, classified as protein-coding or non-coding RNA, and with an associated gene symbol. Lowly expressed genes were further removed using the filterByExpr function with default parameters. Library sizes were normalised using the trimmed mean of M values (TMM) method via normLibSizes. Multidimensional scaling plots were created with limma’s plotMDS function to visualize the gene expression profiles from individual samples (Supplementary Figure 3). A single design matrix was constructed using a means model parametrisation in which each group was defined by the combination of cell type, genotype, and treatment. Differential expression was assessed the enhanced limma-voom pipeline implemented in edgeR’s voomLmFit function (Law, Chen et al. 2014, Baldoni, Chen et al. 2025) while accounting for within-mouse correlation using mouse number as a blocking factor. Sample quality weights were estimated and moderated toward one by setting argument prior.n = 1000 in voomLmFit to stabilise inference given the modest number of replicates per group. Empirical Bayes moderation was applied via eBayes with robust = TRUE to protect against hypervariable genes (Phipson, Lee et al. 2016). Within-cell-type comparisons of Nutlin vs. DMSO in both wild type and p53 KO samples were performed to quantify the p53 transcriptional response. Genes with Benjamini-Hochberg adjusted p-values below 0.05 were considered differentially expressed.

Moderated t-statistics and total degrees of freedom from each Nutlin vs. DMSO comparison were transformed to standard normal z-scores using the function zscoreT from the limma package and compared across cell types and genotypes. The limma function getGeneKEGGLinks was used to match genes and KEGG p53 signaling, apoptosis, cell cycle and cellular senescence pathways (mmu04115, mmu04210, mmu04110, and mmu04218 pathways). The Heatmap function from the ComplexHeatmap (v2.26.1) (Gu, Eils et al. 2016) package was used to visualize the results from both wild type and KO samples for genes associated with the selected KEGG pathway (Figure 3A). Pairwise plots were created to compare Nutlin vs. DMSO z-scores from each cell type for wild type samples (Figure 3C) and p53 KO samples (Supplementary Figure 5).

Self-contained gene set tests were performed using the mroast function from the limma package (Wu, Lim et al. 2010) to assess whether a curated list of genes belonging to p53 signaling, apoptosis, cell cycle arrest, and cellular senescence pathways were differentially expressed as a group upon Nutlin treatment in each cell type. Gene sets for each pathway were curated manually based on well-established pathway members and are presented below.

| Pathway | Genes |
| --- | --- |
| p53 signaling | Cdkn1a, Mdm2, Bbc3, Pmaip1, Gadd45a, Ddb2, Rrm2b, Sesn1, Sesn2, Trp53inp1, Pidd1, Zmat3, Btg2, Perp, Fas, Xpc, Dram1, Igfbp3, Plk2 |
| Apoptosis | Bax, Bak1, Bbc3, Pmaip1, Bcl2, Bcl2l1, Mcl1, Bcl2l11, Bid, Bad, Casp3, Casp7, Casp8, Casp9, Apaf1, Fas, Fadd, Tnfrsf10b, Diablo, Xiap |
| Cell cycle arrest | Cdkn1a, Cdkn2a, Cdkn2b, Rb1, E2f1, E2f2, E2f3, Ccnd1, Ccnd2, Ccne1, Ccne2, Ccna2, Ccnb1, Ccnb2, Cdk1, Cdk2, Cdk4, Cdk6, Cdc25a, Cdc25c, Gadd45a |
| Cellular senescence | Cdkn1a, Cdkn2a, Cdkn2b, Trp53, Rb1, Atm, Atr, Chek1, Chek2, Gadd45a, Glb1, Il6, Cxcl1, Cxcl2, Serpine1, Mmp13, Mmp3, Igfbp7, Tgfb1 |

The mroast function was run with 999,999 rotations and incorporated the blocking factor, intra-block correlation, and sample quality weights estimated by voomLmFit to ensure consistency with the primary differential expression analysis (Figure 3B). Results were visualised using barcode plots generated with limma’s barcodeplot function (Supplementary Figure 4), in which t-statistics from each within-cell-type contrast were ranked and the positions of pathway genes marked to show their relative enrichment among highly up- or down-regulated genes.

### MCL-1 and BCL-XL protein level measurement by flow cytometry

MDFs and BMDMs were plated at 5x10^5^ per well in 6-well plates and left for overnight to adhere. pre-B cells, thymocytes, activated T cells and activated B cells were plated at 1x10^6^ cells per well in 6-well plates. All the cells were treated with either DMSO or nutlin-3a (10 μM) for 24h and incubated in the designated incubator. Q-VD-OPh (25 μM) was added to block cell death during treatment for the suspension cell types. Following the treatment, cells were harvested, fixed and then permeabilised using the eBioscience™ Foxp3/Transcription Factor Staining Buffer Set (Thermo Fisher Scientific, # 00-5523-00) according to the manufacturer’s instructions. Fixed cells were divided into two aliquots, one aliquot was stained with Alexa Fluor 647-conjugated anti-MCL-1 antibody (1:100; WEHI, clone 38/04-19C4-15) and PE-conjugated anti-BCL-XL antibody (1:100; Abcam, #ab208747). The second aliquot was stained with the corresponding Alexa Fluor 647-conjugated IgG isotype control (1:100; WEHI) and PE-conjugated IgG isotype control (1:100; Abcam, #ab209478). Samples were analysed on the Fortessa1 (Becton Dickinson). Data analysis was performed using FlowJo v10 and Prism v9.

### Proteomics sample preparation

MDFs were plated at 5x10^5^ per well in 6-well plates and 2x10^6^ BMDMs were seeded in 9 cm^2^ polystyrene sterile petri dishes (Thermo Fisher, #10002408) and left for overnight to adhere. Thymocytes and activated T cells were plated at 1x10^6^ cells per well in 6-well plates. Human cancer cell lines (A549, RKO, H460 and SJSA-1) were seeded at 1x10^6^ cells per T25 flask (Thermo Fisher, #430168) and allowed to adhere overnight, while DoHH2 cells were plated at 1x10^6^ per well in 6-well plates. All the cells were treated with either DMSO or nutlin-3a (10 μM) for 24 h and incubated in the designated incubator. Q-VD-OPh (25 μM) was added to block cell death during treatment for thymocytes, activated T cells, SJSA-1 and DoHH2. After the treatment, MDFs, BMDMs and human cancer cell lines (A549, RKO, H460 and SJSA-1) were harvested, and the cell pellets were frozen in -80 °C. Thymocytes, activated T cells and DoHH2 were harvested and then resuspended in DPBS containing DAPI (0.5 µg/mL). 1x10^6^ viable cells (DAPI negative) were sorted out by Aria FUVsion (BD Biosciences) for each sample and then frozen in -80 °C. After harvesting, cell pellets were lysed in 100 μL lysis buffer (5% sodium dodecyl sulfate (SDS) (Amresco, #0227), 50 mM TEAB (Thermo Scientific, #90114), and protease inhibitor cocktail (Roche, # 11697498001) 50X in Milli-Q water). Samples were incubated at room temperature for 5 min and mixed at 1000 rpm using a thermomixer for another 5 min. Lysates were then boiled at 95°C for 10 min using a thermomixer at 1000 rpm. Followed by this, samples were sonicated using ultrasonic bath (Fisherbrand, FB15048) for 10 min. To remove trace DNA contamination, Pierce™ universal nuclease (1 μL per sample; Thermo Scientific, #88702) was added, and samples were incubated at 37°C for 15 min. Total protein concentration was then determined using the BCA protein assay kit (Thermo Scientific, #23250) according to the manufacturer’s instructions.

20 μg protein from each sample was bound, cleaned and digested using trypsin (0.5μg per sample; Sigma, ems0004) and lysine C (0.5μg per sample; FUJIFILM Wako Pure Chemical Corporation, #125-05061) on S-Trap^TM^ micro spin column (Protifi, # C002-MICRO-OO80) according to manufacturer’s protocol. As a modification to the manufacturer’s protocol, chloroacetamide (20 mM; Sigma, #C0267) was used as the alkylating reagent instead of MMTS. Peptides were dried in vacuo (Acid Resistant CentriVap Concentrator, Labconco), resuspended in 50 μL 0.1% formic acid (FA) 2% acetonitrile (ACN), and cleaned using GL-Tip ^TM^ SDB column (GL Science, #7820-11200) according to manufacturer’s protocol. Peptides were dried again in vacuo, reconstituted into 10 μL 0.1% FA 2% ACN and transferred into SureSTART™Total Recovery Glass Screw Top Microvials (Thermo, 6PSV9-TR1) for analysis by mass spectrometry.

### Gas phase fractionation data independent acquisition liquid chromatography electrospray tandem mass spectrometry analysis

Data were acquired on an Orbitrap Astral mass spectrometer (Thermo Scientific) coupled to a Vanquish Neo UHPLC system (Thermo Scientific. Peptides were loaded directly onto a 15 cm × 75 µm C18 fused silica column with an integrated emitter tip (Montara Biolabs) at a flow rate of 1.2 µl/min (up to 1500 bar) and separated at 400 nl/min with a 22 min linear gradient from 4% to 34% buffer B, followed by a column wash at 100% buffer B for ∼3 min. Buffer A consisted of water with 0.1% formic acid and buffer B consisted of 80% acetonitrile with 0.1% formic acid.

MS data were acquired in data-independent acquisition (DIA) mode. Full MS survey scans were acquired in the Orbitrap at a resolution of 240,000 over a scan range of 380–980 m/z (AGC target 5.0 × 10⁶, maximum injection time 5 ms), RF lens=50%, data type= profile. DIA scans were acquired in the Astral analyser over a scan range of 145–1450 m/z using 200 user-defined variable-width isolation windows, HCD fragmentation at 25% normalized collision energy, and a cycle time of 0.6 s (AGC target 8.0 × 10⁴, maximum injection time 3 ms), RF lens=40%, data type= centroid).

### Mass spectrometry proteomics data analysis of murine cells

For murine primary cell proteomics, data were processed, searched and quantified with the Spectronaut software package, version 20.2.250922.92449. Data was searched using a library-free approach against the Mouse UniProt database (downloaded 23rd May 2025) and contaminant lists from (Frankenfield, Ni et al. 2022) (Frankenfield, Ni et al. 2022). Raw proteomics data files were searched with the Pulsar tool within Spectronaut, using the BGS Factory settings for peptide and protein identification. Briefly, the digest rule was ‘Trypsin/P’ with a maximum of two missed cleavages and minimum peptide length of 7 amino acids. Carbamidomethyl of cysteine was set as a fixed modification and protein N-terminal acetylation and methionine oxidation as variable modifications. Protein and precursor q-values and PEP cutoffs were all set at default levels.

Downstream analyses of the proteomics data was performed with the packages limpa (v1.3.11) (Li and Smyth 2023, Li, Cobbold et al. 2025) and limma (v3.67.1) (Ritchie, Phipson et al. 2015) in R (v4.5.3; https://www.R-project.org/). First, 226,620 precursor-level intensities from Spectronaut were imported into R with function readSpectronaut. The detection probability curve (DPC) was estimated assuming a complete normal model using the function dpcCN, resulting in estimated model coefficients -5.738482 and 0.958166. The NCBI database (file Mus_musculus.gene_info.gz downloaded from URL https://ftp.ncbi.nlm.nih.gov/gene/DATA/GENE_INFO/Mammalia on 13 February 2026) was used to convert gene aliases to official NCBI gene symbols using limma’s function alias2SymbolUsingNCBI. Precursor ions with no associated gene symbol were removed, resulting in 225,562 precursor ions.

Protein quantification and differential expression analyses were performed for each cell type separately. First, intensities from a total of 9,425 proteins were quantified using the function dpcQuant and the estimated DPC model coefficients. Next, proteins detected in at least three samples were retained for differential testing using function filterByDetection and cyclic loess normalization of protein intensities was performed with limma’s normalizeCyclicLoess function. Multidimensional scaling plots were created with the plotMDSUsingSEs function to visualize the protein intensity profiles from individual samples while accounting for standard errors from the quantification step. Differential expression analysis between DMSO and Nutlin samples was performed using the function dpcDE while accounting for within-mouse correlation using mouse number as a blocking factor, sample quality weights, and protein-level quantification standard errors. Empirical Bayes moderated t-statistics were obtained from limma’s eBayes function. Proteins with Benjamini-Hochberg adjusted p-values below 0.05 were considered differentially expressed.

For quantification used in proteomics ruler pipeline, the proteotypicity filter was set as “none”, protein LFQ method was set to ‘QUANT 2.0 (SN Standard)’, the MS-Level quantity was set to ‘MS2’, major group Top N and minor group Top N were set as ‘False’, and Cross Run Normalization was set to “False”. Default workflow and ID Picker algorithm were used for protein inference and protein Quantity data outputted. Protein quantity data was filtered to remove any proteins identified only from contaminant libraries (Frankenfield, Ni et al. 2022), or that were identified in both Mouse UniProt and contaminants library but were a keratin protein. Data was imported into Perseus version 2.0.10.0 and filtered to include only proteins for which at least one condition had protein quantified in all 4 biological replicates. Protein intensity was normalised and mean copy number per cell estimated using the “proteomic ruler” Perseus plugin as described in (Wiśniewski, Hein et al. 2014). This method assumes equivalent histone content per cell and uses histone signal as an internal standard for data normalisation and estimation of absolute protein abundances per cell. This strategy avoids the error prone steps of cell counting and protein concentration evaluation when estimating absolute abundance and avoids pitfalls from ‘over-normalisation’ when contrasting samples with substantial differences in cell size and protein content as occurs here.

### Mass spectrometry proteomics data analysis of human cell lines

For human cell line proteomics, data were processed, searched and quantified with the Spectronaut software package, version 20.4.260109.92449. Data was searched using a library-free approach against the Human UniProt database (downloaded 19th March 2026) using the BGS Factory settings for peptide and protein identification (as described above).

Downstream analyses were performed in R (v4.6.0) using limpa (v1.4.0) and limma (v3.69.1). The 269,125 precursor-level intensities were imported with readSpectronaut and protein annotation was obtained by mapping UniProt accessions to gene symbols and NCBI gene IDs using a UniProt ID mapping file (downloaded from URL https://ftp.uniprot.org/pub/databases/uniprot/current_release/knowledgebase/idmappi ng/by_organism/HUMAN_9606_idmapping_selected.tab.gz on 15 January 2026) and the Bioconductor annotation package org.Hs.eg.db (v3.23.1). The DPC was estimated assuming a complete normal model using the function dpcCN, resulting in estimated model coefficients -7.24 and 1.09. Precursors detected in fewer than three samples across all 48 samples were removed, as were precursors with no associated gene name, resulting in 250,349 precursors ions.

Protein quantification and differential expression analyses were performed jointly for all cell types. 10,297 proteins were quantified with the function dpcQuant using the DPC estimated by dpcCN. Two samples (one from the DOHH2 cell line and another from the MOLM13 cell line) were identified as clear outliers based on multidimensional scaling plots and excluded from subsequent analyses. Differential expression between Nutlin and DMSO conditions was tested using a design matrix that included a main effect for cell line and cell line-specific treatment contrasts, with heteroscedasticity between groups accounted for via the var.group argument in dpcDE function. Replicate number was included as a blocking factor to account for within-replicate correlation. Empirical Bayes moderated t-statistics were obtained from the eBayes function and proteins with Benjamini-Hochberg adjusted p-values below 0.05 were considered differentially expressed. Gene Ontology and KEGG pathway enrichment analyses were performed using limma’s goana and kegga functions, respectively.

### Retrovirus production, transduction of fetal liver cells (FLCs) and transplantation into lethally irradiated recipient mice

Wild-type human MCL1 and BCL2L1 (BCL-XL) cDNAs cloned into the pMSCV-IRES-GFP (pMIG) retroviral vector were a gift from Prof. Cory (WEHI). Retroviruses were produced by transient transfection of HEK293T cells in 10 cm culture dishes with 3.6 µg of vector DNA and packaging retroviral plasmids Gag-pol (1.2 µg), pENV (1.2 µg) using FuGENE6 (Cat#E2691, Promega) transfection reagent according to the manufacturer’s instructions. Viral supernatants were collected 48-72 h after transfection and passed through a 0.45 µm filter prior to transduction of cells.

Embryonic day E13.5 (E13.5) FLCs were harvested from C57BL/6-Ly5.2 wild-type embryos and transduced with retrovirus supernatants as previously described (Potts, Mizutani et al. 2025). Transduction efficiency was assessed by GFP expression using flow cytometry. Transduced FLCs were transplanted into lethally irradiated C57BL/6-Ly5.1 recipient mice (7–8 weeks old), which had received two doses of 5.5 Gy irradiation administered 3 h apart, as described previously (Potts, Mizutani et al. 2025).

### Cut and Run assay

MDFs were plated at 5x10^5^ per well in 6-well plates and left for overnight to adhere. Thymocytes were seeded at 5x10^5^ per well in 24-well plates immediately prior to treatment. Cells were treated with either DMSO or nutlin-3a (10 μM) for 6h. To prevent apoptosis during treatment, thymocytes were cultured in the presence of the pan-caspase inhibitor Q-VD-OPh (25 μM). Following treatment, cells were harvested, washed once with DPBS at room temperature, and resuspended in 500 μL ice-cold NEI buffer (20 mM HEPES-KOH, pH 7.9, 10 mM KCl, 0.5 mM spermidine, 0.1% Triton X-100, 20% glycerol, and protease inhibitor cocktail) with gentle vortexing. Cell suspensions were incubated on ice for 10 min to permeabilize cell membrane and release nuclei. Nuclei were then pelleted by centrifugation at 1,300 × g for 4 min at 4°C. After removal of the supernatant, the nuclei were gently resuspended in 100 μL Wash Buffer supplied with the CUTANA™ ChIC/CUT&RUN Kit v6 (EpiCypher, #14-1408). CUT&RUN samples were subsequently prepared according to the manufacturer’s instructions. For each sample, 1 μg of anti-p53 antibody (Leica Biosystems, # NCL-L-p53-CM5p) or rabbit IgG control antibody supplied with the kit was used. CUT&RUN DNA was used for preparing Illumina sequencing libraries using the CUTANA™ CUT&RUN Library Prep Kit (EpiCypher, #14-1001) according to the manufacturer’s instructions. Libraries were then sequenced on an Illumina NextSeq platform, 10 million reads per sample were collected.

### CUT&Run data analysis

CUT&RUN samples were collected from thymocytes and MDFs across two genotypes (WT and p53 KO) and two treatment conditions (Nutlin and DMSO), with IgG used as a negative control, yielding 16 samples in total (8 per cell type) without biological replicates. Raw paired-end FASTQ files were aligned to the mouse reference genome mm39 using Rubread with arguments sortReadsByCoordinates = TRUE and maxFragLength = 2000.

Following alignment, read filtering was performed with samtools (v1.21) (Danecek, Bonfield et al. 2021) to remove unmapped reads, reads with unmapped mates, non-primary alignments, reads failing platform quality checks, reads with a mapping quality below 30, and improperly paired reads. Orphan reads and read pairs mapping to different chromosomes were removed using samtools fixmate. Duplicate reads were marked using Picard MarkDuplicates (v3.3.0; https://broadinstitute.github.io/picard) and then removed using samtools, yielding the final filtered, deduplicated BAM files used for all downstream analyses.

BigWig files were generated from the deduplicated BAM files using bamCoverage from the deepTools suite (v3.5.1) (Ramírez, Ryan et al. 2016), with RPGC normalisation (1x genomic coverage) and a bin size of 10 bp relative to the reference autosome. The effective genome size was set to 2,654,605,538 bp, corresponding to the mappable mm39 mouse genome. Peak calling was performed using MACS3 (v3.0.1) (Zhang, Liu et al. 2008) on each of the 8 p53 CUT&RUN samples from thymocytes and MDFs, with the matched IgG sample from the same cell type, genotype, and treatment condition used as the control. MACS3 was run with options -f BAMPE -g 2654605538 -q 0.01 --call-summits --bdg, yielding one set of peaks per sample covering all combinations of genotype (WT and KO) and treatment (Nutlin and DMSO) for each cell type.

Signal enrichment heatmaps were generated for each cell type using the deepTools suite. A cell type-specific consensus peak set was derived by merging the WT DMSO and WT Nutlin p53 narrowPeak files using bedtools merge (v2.31.1) (Quinlan and Hall 2010). Signal matrices were computed using computeMatrix in reference-point mode, centred on the peak midpoint and extending 2 kb in both directions with a bin size of 10 bp, separately for each of the p53 antibody samples WT Nutlin, WT DMSO, KO Nutlin, and KO DMSO. Heatmaps were generated using plotHeatmap with peaks sorted in descending order of mean signal intensity (Supplementary Figures 11A–B).

Coverage plots were generated using the plotgardener package (v1.16.0) (Kramer, Davis et al. 2022), with bigWig files imported using the import.bw function from the rtracklayer package (v1.70.1) (Lawrence, Gentleman et al. 2009). For each gene of interest, genomic coordinates were retrieved from the Bioconductor annotation package TxDb.Mmusculus.UCSC.mm39.knownGene package (v3.22.0) and signal tracks were plotted for all WT p53 samples from for each cell type) (Supplementary Figures 11C–D).

### Lentivirus production and CRISPR/Cas9 library screening

Lentiviruses were produced by transient transfection of HEK293T cells cultured in ten 10-cm tissue culture dishes. Each dish was transfected with 1.5 µg of YUSA sgRNA library DNA together with the lentiviral packaging plasmids pMDLg (1.5 µg), pRSV-REV (1.5 µg), and VSV-G (1 µg) using FuGENE6 Transfection Reagent (Promega, #E2691) according to the manufacturer’s instructions. Viral supernatants from all dishes were collected 48–72 h after transfection, pooled, and passed through a 0.45 µm filter prior to transduction of target cells.

Cas9-expressing MEFs were seeded in twenty 10-cm tissue culture dishes at a density of 3.5 × 10^6 cells per dish and allowed to adhere overnight. Cells were then transduced with lentiviral supernatants for 48 h, achieving >60% transduction efficiency as determined by BFP expression. Library coverage of approximately 300× per sgRNA was maintained throughout the experiment, with ∼99.6% representation of the sgRNA library detected in the input population. Cells were subsequently treated with Nutlin-3a (10 μM) for 24 h. Plasma membranes were permeabilised in 0.025% digitonin in MELB buffer (10 mM HEPES-KOH, pH 7.5, 25 mM KCl, 10 mM MgCl₂, and 10 μM DTT) for 10 min on ice, and cells were stained with PE-conjugated anti-BCL-XL antibody (1:100 dilution in MELB buffer) for 30 min on ice. Based on PE fluorescence intensity, cells were sorted into three populations: the dimmest 5%, the brightest 5%, and the remaining 90% of cells. An aliquot (12×10^6 cells) of the unsorted population was collected as the input control. Genomic DNA was extracted from each population using the DNeasy Blood & Tissue Kit (QIAGEN, #69504) according to the manufacturer’s instructions. sgRNA sequences were amplified by PCR and sequenced as previously described (La Marca, Aubrey et al. 2024). Libraries were pooled and sequenced using an Illumina 100 million-read kit, generating approximately 20–25 million reads per sample. sgRNAs enriched in the dimmest or brightest populations were identified by comparison with sgRNA representation in the input population using MAGeCK.

## Supplementary Figure Legends

**Supplementary Figure 1:** F**old change of GFP (*p21-IRES-GFP* reporter) and tdTomato (*Puma-tdTomato* reporter) expression upon p53 activation.**

MDFs (**A–B**), BMDMs (**C–D**), activated T cells (**E–F**), and activated B cells (**G–H**) derived from *p21-IRES-GFP^KI/+^;Puma-tdTomato^KI/+^* double reporter mice were treated *in vitro* with DMSO or 10 µM nutlin-3a for the indicated time points. For activated T cells and activated B cells, the caspase inhibitor Q-VD-OPh was included to prevent cell demolition due to apoptosis. Cells were analysed for GFP (*p21-IRES-GFP* reporter) and tdTomato (*Puma-tdTomato* reporter) expression by flow cytometry. Data are presented as fold change relative to DMSO treated control cells. Data are presented as mean ± SD (n = 3 biological replicates). Statistical significance was determined using two-way ANOVA with Dunnett’s multiple comparisons test (P < 0.05; P < 0.01; P < 0.001; P < 0.0001; ns, not significant).

**Supplementary Figure 2: Loss of *P21*(*Cdkn1a*) attenuates p53-induced cell cycle arrest but does not impact apoptosis.**

**A, C, E** and **G**: MDFs (**A**), BMDMs (**C**), activated T cells (**E**) and activated B cells (**G**) from C57BL/6 mice (wt) and *P21*(*Cdkn1a)*^-/-^ mice were treated with 10 µM nutlin-3a for the indicated time points (MDFs and BMDMs were analysed after 24 h, 48 h and 72 h, activated T cells and activated B cells were analysed after 24 h and 48 h). Apoptosis was assessed by Annexin-V plus DAPI staining followed by flow cytometry (Annexin V-DAPI- cells were deemed live cells). Data are presented as mean ± SD (n = 3 biological replicates).

**B, D, F** and **H**: Cell cycle distribution of MDFs (**B**), BMDMs (**D**), activated T cells (**F**) and activated B cells (**H**) after 24 h treatment with 10 µM nutlin-3a with BrdU added to the cultures during the final 4h. Cell cycle distribution was determined by staining for BrdU incorporation and DNA content (7-AAD) and cells were divided into the following stages of the cell cycle: sub-G1 (apoptotic), G0/G1, S, or G2/M. For activated T cells and activated B cells, dead cells were excluded, and cell cycle analysis was performed on viable cells only. Data are presented as mean ± SD (n = 3 biological replicates).

**Supplementary Figure 3: MDS plots of RNA-seq data highlight transcriptomic differences between cell types.**

Multidimensional scaling (MDS) plots of RNA-seq data from the indicated *Trp53⁻/⁻* (**A**) and wild-type cells (**B**), illustrating transcriptomic differences across different cell types.

**Supplementary Figure 4: Gene set enrichment analysis across p53-related pathways**

**A-D:** Barcode plots showing self-contained gene set testing for each cell type across p53-related pathways: p53 pathway (**A**), apoptosis pathway (**B**), cell cycle pathway (**C**) and cell senescence pathway (**D**).

**Supplementary Figure 5: Pairwise comparison of transcriptional responses in *Trp53⁻/⁻* cells**

Pairwise plots showing Z-scores derived from t-statistics comparing nutlin-3a treatment versus DMSO treatment within each cell type from *Trp53⁻/⁻* mice.

**Supplementary Figure 6: Flow cytometry histograms showing changes in BCL-XL and MCL-1 protein levels following p53 activation across different primary murine cell types**

MDFs, BMDMs, activated T cells, activated B cells, thymocytes, and pre-B cells from wild-type mice (C57BL/6) were treated for 24 h with DMSO or 10 µM nutlin-3a in the presence of 25 µM QVD-OPH. Protein levels of MCL-1 (**A**) and BCL-XL (**C**) were assessed by intra-cellular staining and flow cytometry using A647-conjugated MCL-1 or PE-conjugated BCL-XL antibodies. Corresponding Ig isotype matched antibodiess (**B**, **D**) were used as controls. Data are representative of three independent experiments using cells derived from n=3 mice.

**Supplementary Figure 7: Blocking apoptosis does not impact changes in BCL-XL or MCL-1 levels after p53 activation**

**A:** Schematic illustrating the generation of *Bak⁻/⁻Bax⁻/⁻* thymocytes, activated T cells, and activated B cells. Foetal liver cells (a rich source of haematopoietic stem/progenitor cells (HSPCs)) from *Bak⁻/⁻Bax⁻/⁻* C57BL/6-Ly5.2 E14.5 embryos, generated by homotypic intercrosses of *Bak⁻/⁻Bax⁺/⁻* mice, or from control wild-type C57BL/6-Ly5.2 embryos were transplanted into lethally irradiated C57BL/6-Ly5.1 recipient mice. Reconstitution efficiency (>95% Ly5.2⁺ cells) was confirmed in peripheral blood of recipient mice at 8-10 post-transplantation by flow cytometric analysis after staining with antibodies against Ly5.1 (identifying host derived cells) and Ly5.2 (identifying donor derived cells).

**B:** Thymocytes from the mice described in A were treated *in vitro* with 10 µM nutlin-3a for 12 h and 24 h. Apoptosis was assessed by staining with Annexin-V plus DAPI followed by flow cytometry (Annexin V-DAPI- cells were deemed live cells). Data are presented as mean ± SD (n = 3 biological replicates).

**C** and **D:** Summary of intracellular flow cytometric analysis of MCL-1 (**C**) and BCL-XL

(**D**) protein levels in thymocytes from mice reconstituted with a wild-type or *Bak⁻/⁻Bax⁻/⁻* haematopoietic system following 24 h treatment with DMSO or 10 µM nutlin-3a in the presence of 25 µM QVD-OPH. Protein levels of MCL-1 and BCL-XL were assessed by intra-cellular staining and flow cytometry using A647-conjugated MCL-1 or PE-conjugated BCL-XL antibodies. Corresponding Ig isotype matched antibodies were used as controls. MCL-1 and BCL-XL protein abundance was quantified by median fluorescence intensity (MFI) following subtraction of the corresponding Ig isotype matched control antibody staining and normalisation to DMSO treated control cells. Statistical significance was determined using two-way ANOVA with Dunnett’s multiple comparisons test (P < 0.05; P < 0.01; P < 0.001; P < 0.0001; ns, not significant). Data are presented as mean ± SD (n = 3 biological replicates).

**E**: Activated T cells generated from mice reconstituted with a wild-type or *Bak⁻/⁻Bax⁻/⁻* haematopoietic system were treated *in vitro* with 10 µM nutlin-3a for 24 h and 48 h. Apoptosis was analysed as described in panel **A**. Data are presented as mean ± SD (n = 3 biological replicates).

**F**: Cell cycle distribution of activated T cells generated from mice reconstituted with a wild-type or *Bak⁻/⁻Bax⁻/⁻* haematopoietic system were treated *in vitro* with 10 µM nutlin-3a for 72 h with BrdU added for the final ***XXX*** h of culture. Cell cycle distribution was determined by staining for BrdU incorporation and DNA content (7-AAD) and cells were divided into the following stages of the cell cycle: sub-G1 (apoptotic), G0/G1, S, or G2/M. Data are presented as mean ± SD (n = 3 biological replicates).

**G** and **H:** Summary of intracellular flow cytometric analysis of MCL-1 (**G**) and BCL-XL

(**H**) protein levels in activated T cells generated from mice reconstituted with a wild-type or *Bak⁻/⁻Bax⁻/⁻* haematopoietic system following 24 h treatment with DMSO or 10 µM nutlin-3a in the presence of 25 µM QVD-OPH. Protein abundance was quantified as described in C and D. Data are presented as mean ± SD (n = 3 biological replicates).

**I**: Activated B cells generated from mice reconstituted with a wild-type or *Bak⁻/⁻Bax⁻/⁻* haematopoietic system were treated *in vitro* with 10 µM nutlin-3a for 24 h and 48 h. Apoptosis was analysed as described in (**A**). Data are presented as mean ± SD (n = 3 biological replicates).

**J**: Cell cycle distribution of activated B cells generated from mice reconstituted with a wild-type or *Bak⁻/⁻Bax⁻/⁻* haematopoietic system were treated *in vitro* with 10 µM nutlin-3a for 48 h with BrdU added during the final 4h of culture. Cell cycle distribution was determined as described in (**F**). Data are presented as mean ± SD (n = 3 biological replicates).

**K** and **L:** Summary of intracellular flow cytometric analysis of MCL-1 (**G**) and BCL-XL

(**H**) protein levels in activated B cells generated from mice reconstituted with a wild-type or *Bak⁻/⁻Bax⁻/⁻* haematopoietic system following 24 h treatment with DMSO or 10 µM nutlin-3a in the presence of 25 µM QVD-OPH. MCL-1 and BCL-XL protein abundance was quantified as described in panels C and D. Data are presented as mean ± SD (n = 3 biological replicates).

**Supplementary Figure 8: Pharmacological inhibition of BCL-XL and/or MCL-1 converts p53-induced cell cycle arrest to apoptosis across multiple primary murine cell types.**

**A-D:** MDFs (**A**), BMDMs (**B**), activated T (**C**) and activated B (**D**) were treated for 24 h with 1 µM MCL-1 inhibitor (S63845), 1 µM BCL-XL inhibitor (A-1331852), 10 µM Nutlin-3a or the indicated combinations of agents with BrdU added for the final 4 h of culture. Cell cycle distribution was analysed by staining for BrdU incorporation and DNA content (7-AAD). Cells were classified as sub-G1 (apoptotic), G0/G1, S, or G2/M. Data are representative of n = 3 independent mice.

**Supplementary Figure 9: BCL-XL and/or MCL-1 upregulation after p53 activation in human cancer cell lines resistant to apoptosis.**

**A** and **B**: Summary of intracellular flow cytometric analysis of MCL-1 (**A**) and BCL-XL (*BCL2L1*) (**B**) protein levels in A549, RKO, H460, SJSA-1 and DOHH2 cells following 24 h treatment with DMSO or nutlin-3a (10 µM) in the presence of 25 µM QVD-OPH. MCL-1 and BCL-XL protein levels were assessed by intra-cellular staining and flow cytometry using A647-conjugated MCL-1 or PE-conjugated BCL-XL antibodies. Corresponding Ig isotype matched antibodies were used as controls. MCL-1 and BCL-XL protein abundance was quantified by median fluorescence intensity (MFI) following subtraction of the corresponding Ig isotype matched control antibody staining and normalisation to DMSO treated control cells. Statistical significance was determined using two-way ANOVA with Dunnett’s multiple comparisons test (P < 0.05; P < 0.01; P < 0.001; P < 0.0001; ns, not significant). Data are presented as mean ± SD (n = 3 biological replicates).

**Supplementary Figure 10: Genome-wide CRISPR/Cas9 library gene deletion screen identifies that p53 is essential for upregulation of BCL-XL after treatment with nutlin-3a.**

**A:** Schematic of the genome-wide CRISPR/Cas9 gene deletion screen. Primary MEFs were derived from Cas9^KI/+^ C57BL/6 E14.5 embryos generated by crossing Cas9^KI/KI^ CS&BL/6 mice with wild-type C57BL/6 mice. Cells were transduced with a lentiviral sgRNA library and, after 48 h, treated for 24 h with 10 µM nutlin-3a. Cells were then permeabilized and stained with a PE-conjugated BCL-XL antibody, and cells with low BCL-XL expression were sorted and collected. Enriched sgRNAs were identified by DNA sequencing.

**B:** sgRNAs targeting *p53* were significantly enriched in the BCL-XL-low cell population compared with the input library.

**C:** Flow cytometry analysis showing that BCL-XL upregulation following p53 activation is p53-dependent. MDFs from wild-type and *p53⁻/⁻* mice (C57BL/6 background) were treated for 24 h with DMSO or 10 µM nutlin-3a, and BCL-XL protein levels were assessed by immunostaining and flow cytometry using a PE-conjugated BCL-XL antibody.

**Supplementary Figure 11: p53 binds to *Bcl2l1* loci at the distal region in MDFs and thymocytes**

**A and B:** Heatmap from global analysis of p53 CUT&RUN data showing the read coverage 2kbp up- and downstream of enriched peaks found genome-wide in MDFs(**A**) and thymocytes (**B**) derived from wild-type and Trp53^-/-^ were treated with DMSO or 10 µM nutlin-3a for 6h. Peaks were called with MACS2.

**C:** p53 antibody CUT&RUN tracks for the known TRP53 target genes *Cdkn1a/P21*, *Bbc3/Puma* and *Mdm2*. MDFs and thymocytes derived from wild-type were treated with DMSO or 10 µM nutlin-3a for 6h. Peaks were called with MACS2.

**D:** p53 antibody CUT&RUN tracks for *Bcl2l1*.

**Supplementary Figure 12: Levels of p53 protein after treatment with nutlin-3a in MDFs, BMDMs, activated T cells and thymocytes.**

**A** and **B:** Copy numbers of p53 (**A**) and MDM2 proteins (**B**) per million cells in MDFs, BMDMs, activated T cells, and thymocytes from C57BL/6 wild-type mice before and after 24 h after treatment with nutlin-3a were quantified by mass-spectrometry using the histone ruler method (copy number per million cells).

