## Supplementary figures for "Pre-existing levels of pro-survival proteins and induction of BCL-XL dictate cell fate after p53 activation"

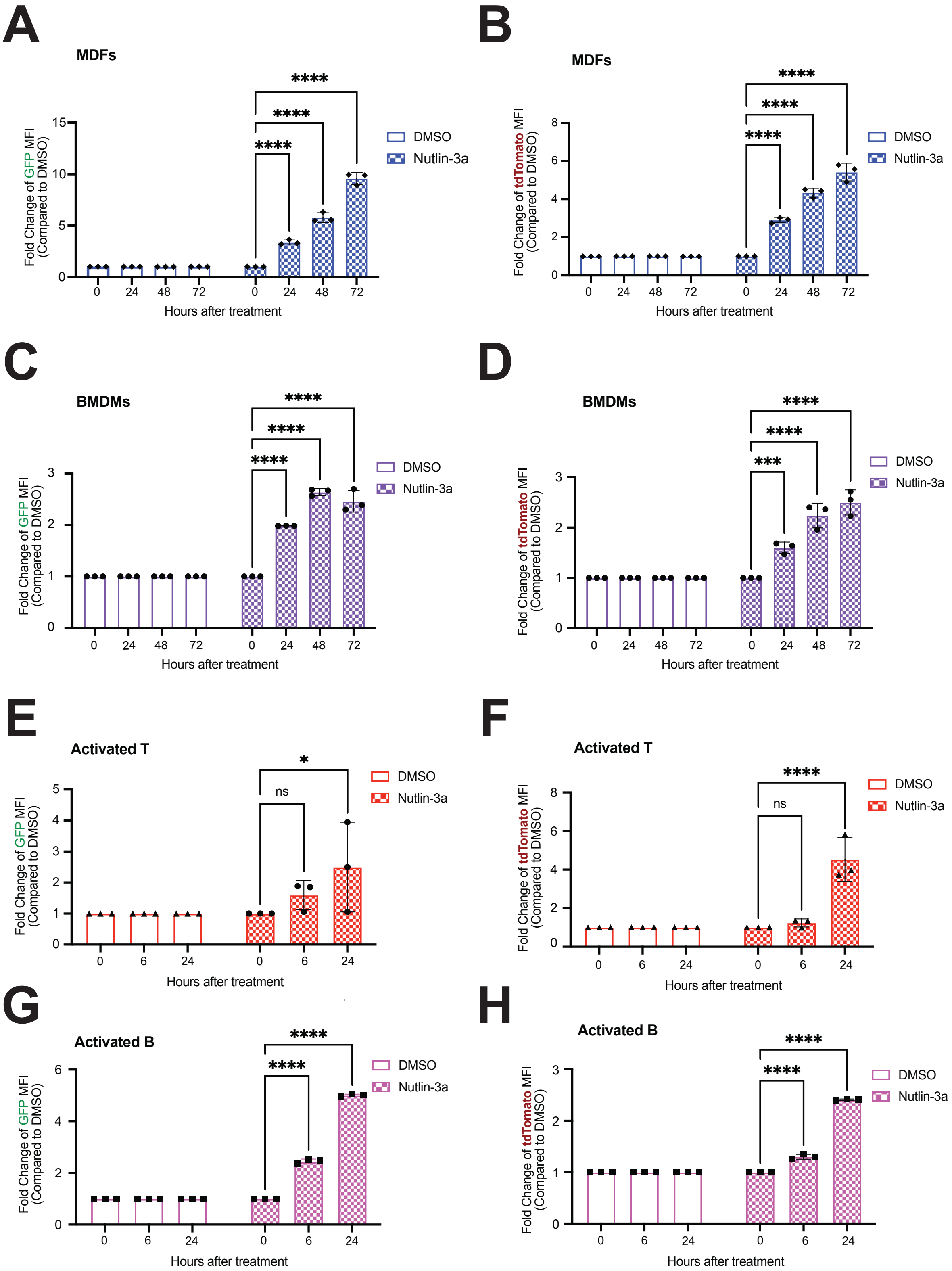

A
MDFs
Fold Change of GFP MFI (Compared to DMSO)
Hours after treatment
DMSO
Nutlin-3a
B
MDFs
Fold Change of tdTomato MFI (Compared to DMSO)
C
BMDMs
D
BMDMs
E
Activated T
ns
F
Activated T
G
Activated B
H
Activated B

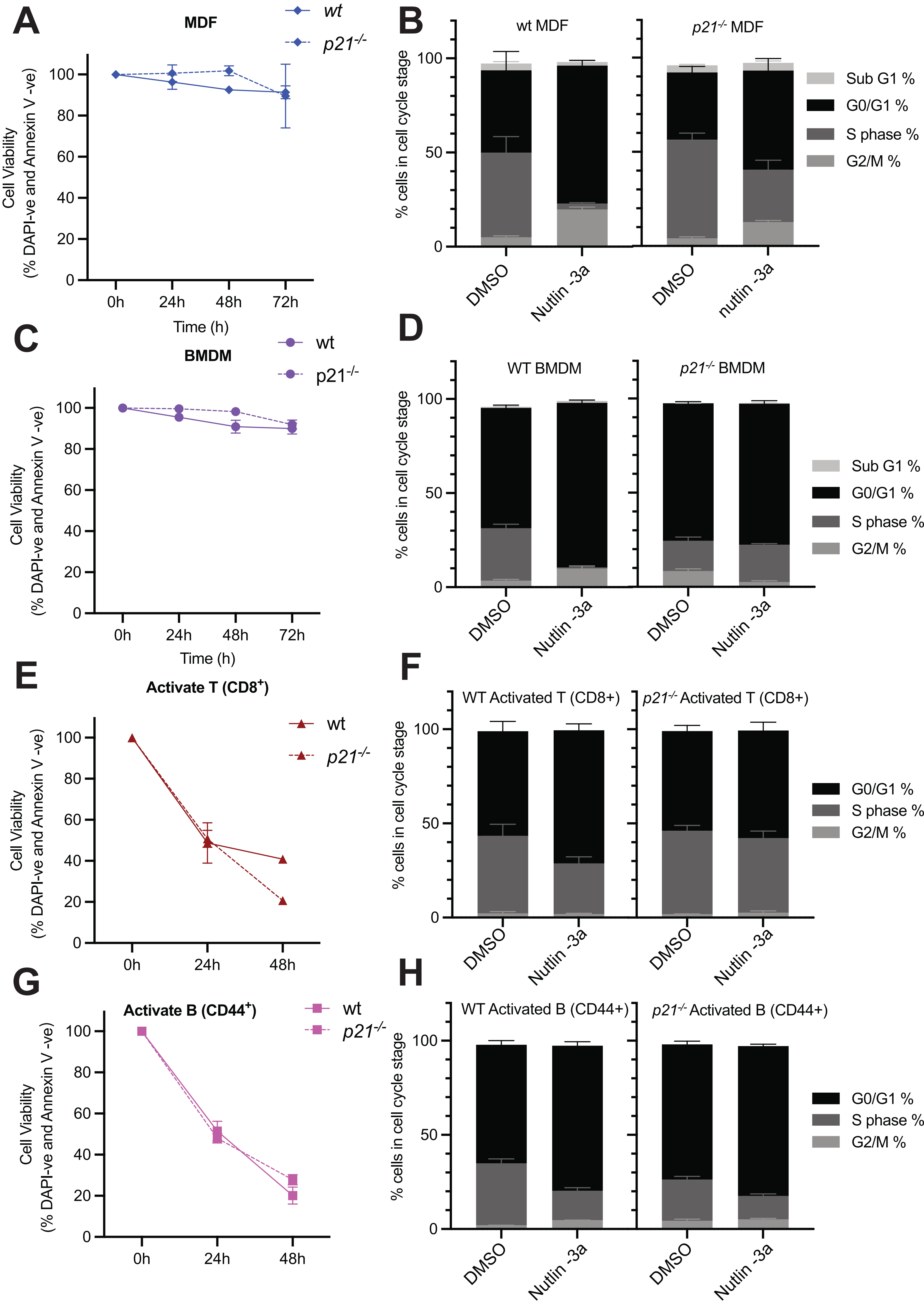

A
MDF
wt
p21-/-
Cell Viability
(% DAPI-ve and Annexin V -ve)
Time (h)
0h
24h
48h
72h
B
wt MDF
p21-/- MDF
% cells in cell cycle stage
DMSO
Nutlin -3a
nutlin -3a
Sub G1 %
G0/G1 %
S phase %
G2/M %
C
BMDM
D
WT BMDM
p21-/- BMDM
E
Activate T (CD8+)
F
WT Activated T (CD8+)
p21-/- Activated T (CD8+)
G
Activate B (CD44+)
H
WT Activated B (CD44+)
p21-/- Activated B (CD44+)

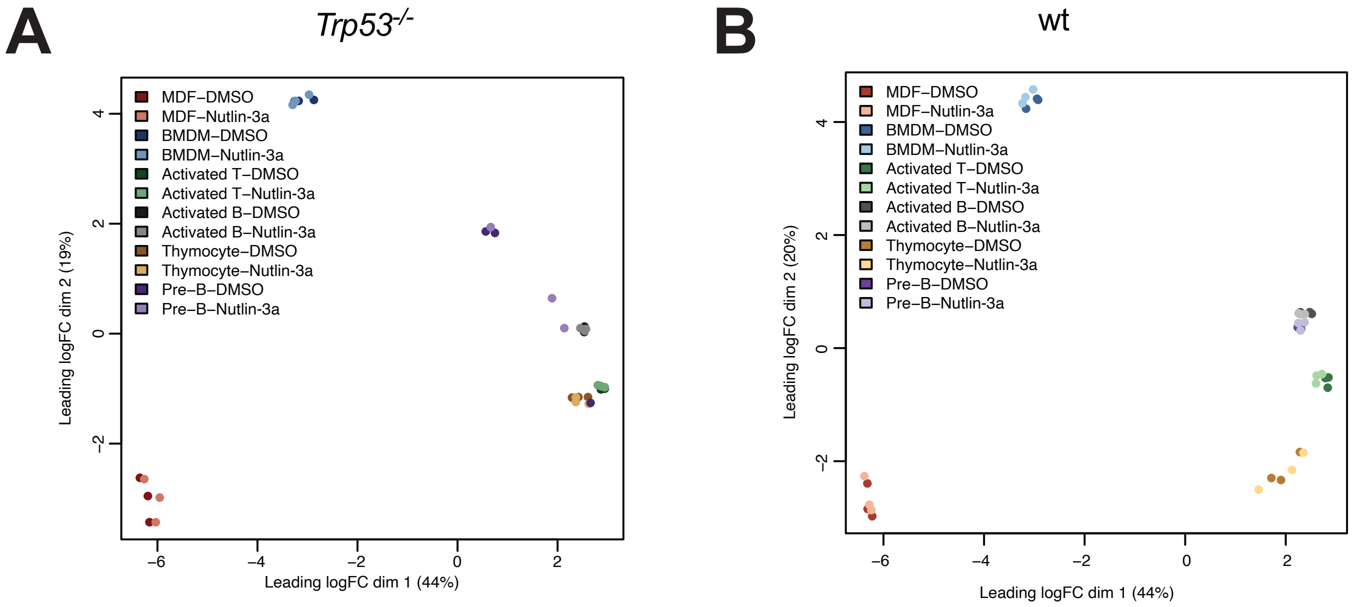
A
Trp53-/-
MDF–DMSO
MDF–Nutlin-3a
BMDM–DMSO
BMDM–Nutlin-3a
Activated T–DMSO
Activated T–Nutlin-3a
Activated B–DMSO
Activated B–Nutlin-3a
Thymocyte–DMSO
Thymocyte–Nutlin-3a
Pre–B–DMSO
Pre–B–Nutlin-3a
Leading logFC dim 2 (19%)
Leading logFC dim 1 (44%)
B
wt
MDF–DMSO
MDF–Nutlin-3a
BMDM–DMSO
BMDM–Nutlin-3a
Activated T–DMSO
Activated T–Nutlin-3a
Activated B–DMSO
Activated B–Nutlin-3a
Thymocyte–DMSO
Thymocyte–Nutlin-3a
Pre–B–DMSO
Pre–B–Nutlin-3a
Leading logFC dim 2 (20%)
Leading logFC dim 1 (44%)

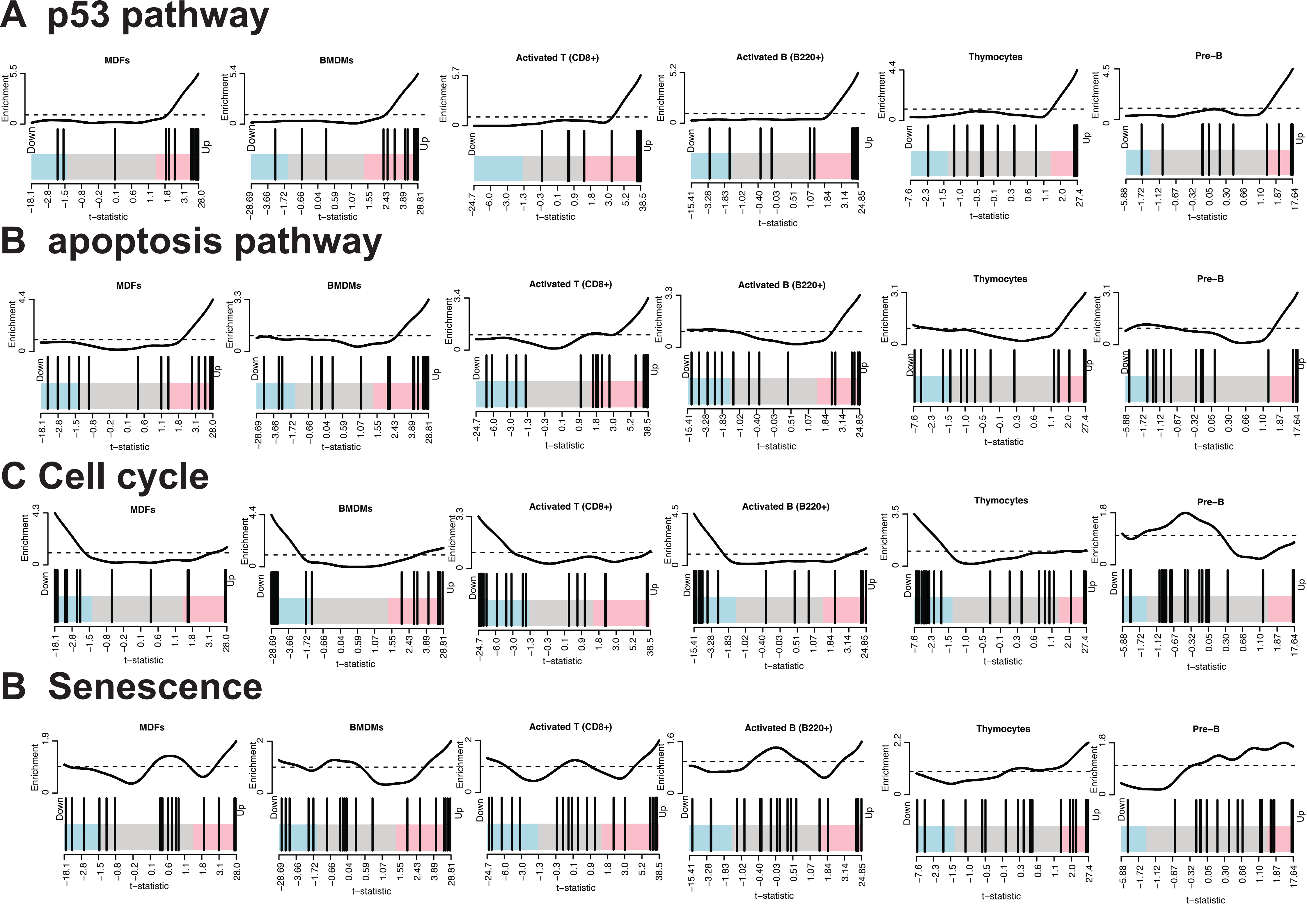
A p53 pathway
B apoptosis pathway
C Cell cycle
B Senescence
MDFs
BMDMs
Activated T (CD8+)
Activated B (B220+)
Thymocytes
Pre−B
Enrichment
Down
Up
t−statistic

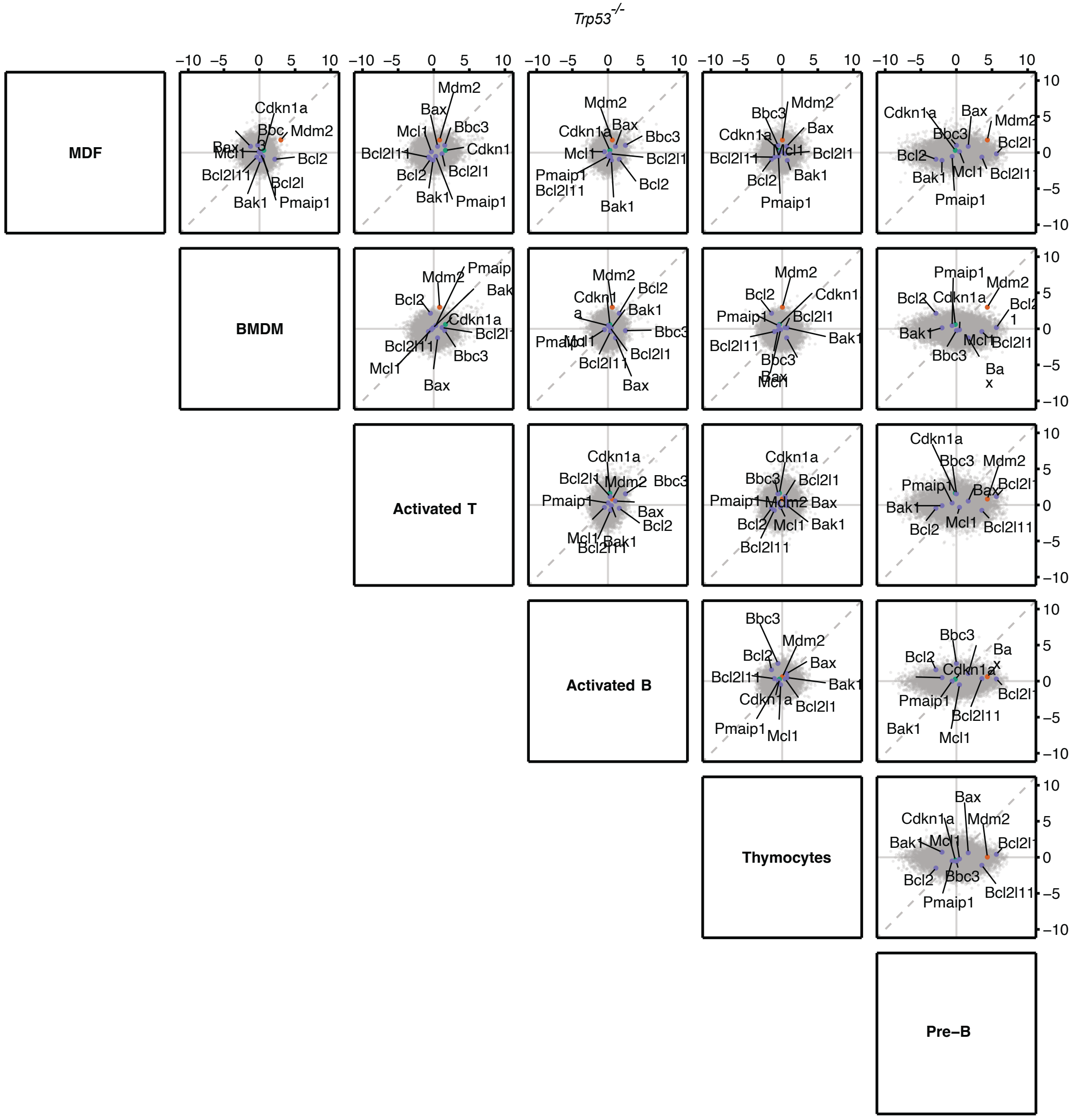

Trp53-/-
MDF
BMDM
Activated T
Activated B
Thymocytes
Pre–B

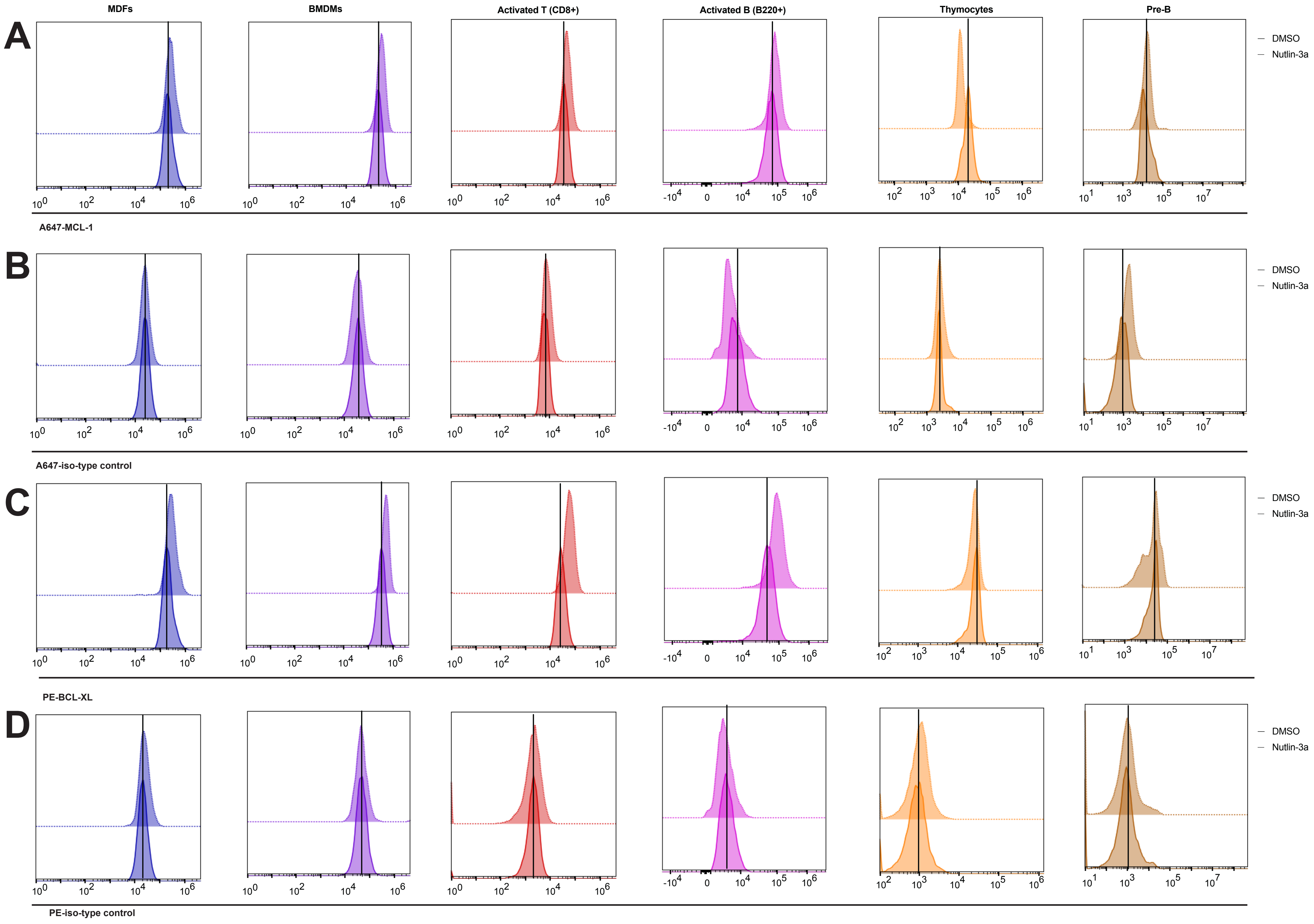
MDFs
BMDMs
Activated T (CD8+)
Activated B (B220+)
Thymocytes
Pre-B
A
B
C
D
DMSO
Nutlin-3a
A647-MCL-1
A647-iso-type control
PE-BCL-XL
PE-iso-type control
$10^0$
$10^2$
$10^4$
$10^6$
$-10^4$
0
$10^5$
$10^3$
$10^1$
$10^7$

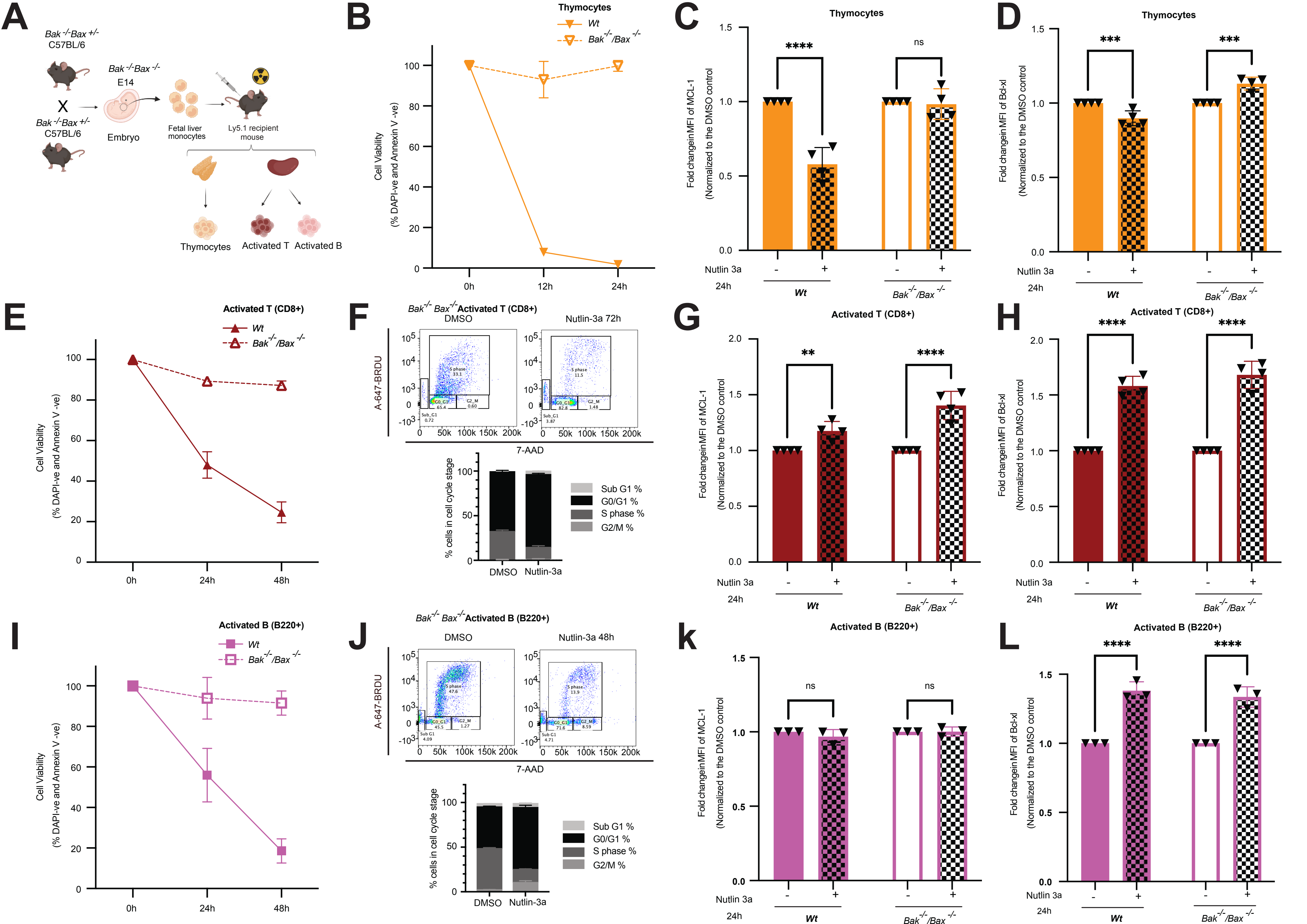

A
Bak -/-Bax +/- C57BL/6
X
Bak -/-Bax +/- C57BL/6
Bak -/-Bax -/- E14
Embryo
Fetal liver monocytes
Ly5.1 recipient mouse
Thymocytes
Activated T
Activated B
B
Thymocytes
Wt
Bak-/-/Bax-/-
Cell Viability (% DAPI-ve and Annexin V -ve)
0h
12h
24h
C
Thymocytes
Fold changein MFI of MCL-1 (Normalized to the DMSO control)
****
ns
Nutlin 3a
24h
Wt
Bak-/-/Bax-/-
D
Thymocytes
Fold changein MFI of Bcl-xl (Normalized to the DMSO control)
***
***
E
Activated T (CD8+)
Wt
Bak-/-/Bax-/-
Cell Viability (% DAPI-ve and Annexin V -ve)
0h
24h
48h
F
Bak-/- Bax-/- Activated T (CD8+)
DMSO
Nutlin-3a 72h
A-647-BRDU
7-AAD
S phase
G0_G1
G2_M
Sub_G1
% cells in cell cycle stage
Sub G1 %
G0/G1 %
S phase %
G2/M %
DMSO
Nutlin-3a
G
Activated T (CD8+)
Fold changein MFI of MCL-1 (Normalized to the DMSO control)
**
****
H
Activated T (CD8+)
Fold changein MFI of Bcl-xl (Normalized to the DMSO control)
****
****
I
Activated B (B220+)
Wt
Bak-/-/Bax-/-
Cell Viability (% DAPI-ve and Annexin V -ve)
J
Bak-/- Bax-/- Activated B (B220+)
DMSO
Nutlin-3a 48h
A-647-BRDU
7-AAD
% cells in cell cycle stage
Sub G1 %
G0/G1 %
S phase %
G2/M %
k
Activated B (B220+)
Fold changein MFI of MCL-1 (Normalized to the DMSO control)
ns
ns
L
Activated B (B220+)
Fold changein MFI of Bcl-xl (Normalized to the DMSO control)
****
****
Nutlin 3a
24h
Wt
Bak-/-/Bax-/-

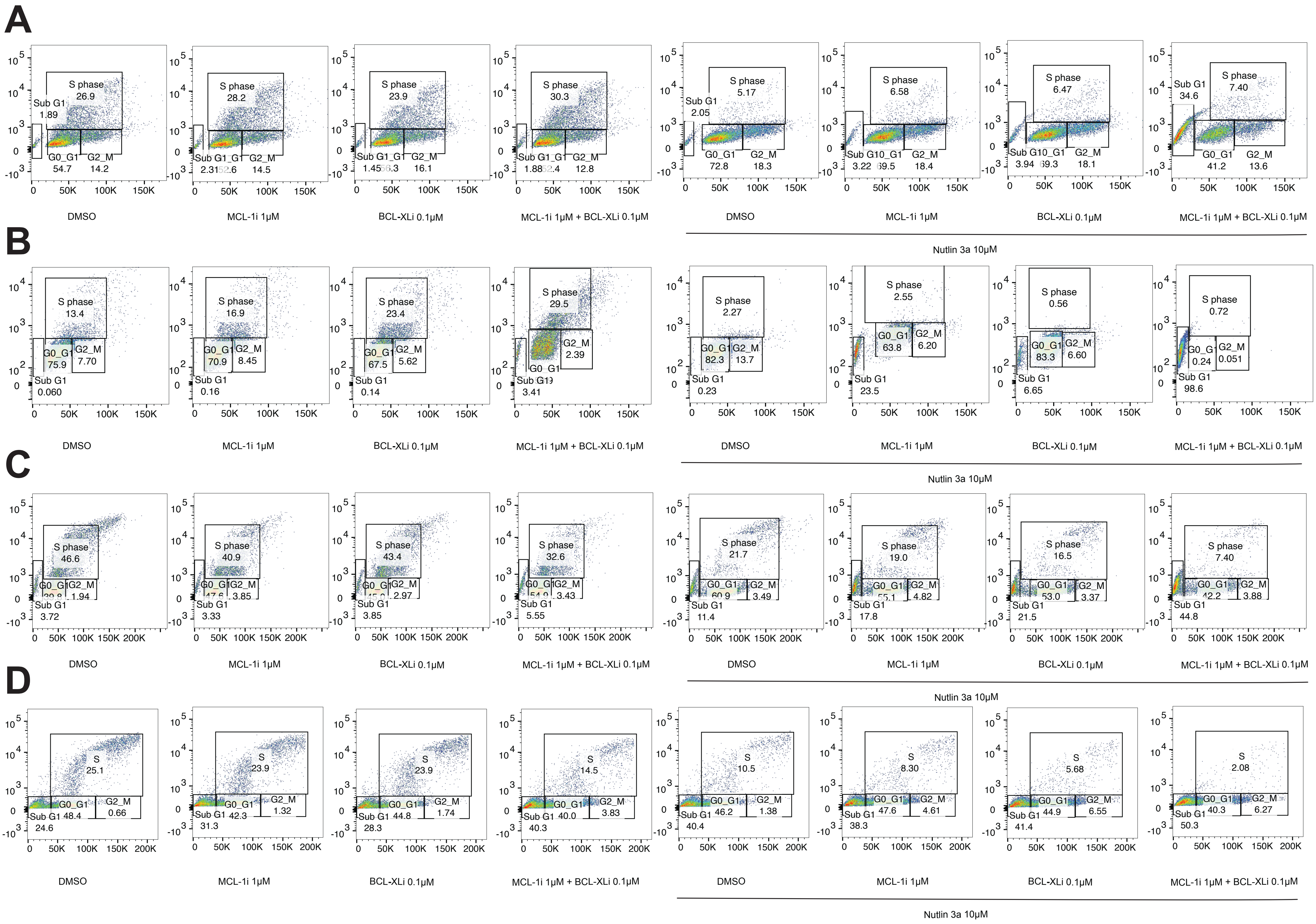

A
S phase 26.9
Sub G1 1.89
G0_G1 54.7
G2_M 14.2
S phase 28.2
G2_M 14.5
S phase 23.9
G2_M 16.1
S phase 30.3
G2_M 12.8
S phase 5.17
Sub G1 2.05
G0_G1 72.8
G2_M 18.3
S phase 6.58
G2_M 18.4
S phase 6.47
G2_M 18.1
S phase 7.40
Sub G1 34.6
G0_G1 41.2
G2_M 13.6
DMSO
MCL-1i 1μM
BCL-XLi 0.1μM
MCL-1i 1μM + BCL-XLi 0.1μM
Nutlin 3a 10μM
B
S phase 13.4
G0_G1 75.9
G2_M 7.70
Sub G1 0.060
S phase 16.9
G0_G1 70.9
G2_M 8.45
Sub G1 0.16
S phase 23.4
G0_G1 67.5
G2_M 5.62
Sub G1 0.14
S phase 29.5
G2_M 2.39
Sub G1 3.41
S phase 2.27
G0_G1 82.3
G2_M 13.7
Sub G1 0.23
S phase 2.55
G0_G1 63.8
G2_M 6.20
Sub G1 23.5
S phase 0.56
G0_G1 83.3
G2_M 6.60
Sub G1 6.65
S phase 0.72
G0_G1 0.24
G2_M 0.051
Sub G1 98.6
C
S phase 46.6
G2_M 1.94
Sub G1 3.72
S phase 40.9
G2_M 3.85
Sub G1 3.33
S phase 43.4
G2_M 2.97
Sub G1 3.85
S phase 32.6
G2_M 3.43
Sub G1 5.55
S phase 21.7
G0_G1 60.9
G2_M 3.49
Sub G1 11.4
S phase 19.0
G2_M 4.82
Sub G1 17.8
S phase 16.5
G0_G1 53.0
G2_M 3.37
Sub G1 21.5
S phase 7.40
G0_G1 42.2
G2_M 3.88
Sub G1 44.8
D
S 25.1
G0_G1 48.4
G2_M 0.66
Sub G1 24.6
S 23.9
G0_G1 42.3
G2_M 1.32
Sub G1 31.3
S 23.9
G0_G1 44.8
G2_M 1.74
Sub G1 28.3
S 14.5
G0_G1 40.0
G2_M 3.83
Sub G1 40.3
S 10.5
G0_G1 46.2
G2_M 1.38
Sub G1 40.4
S 8.30
G0_G1 47.6
G2_M 4.61
Sub G1 38.3
S 5.68
G0_G1 44.9
G2_M 6.55
Sub G1 41.4
S 2.08
G0_G1 40.3
G2_M 6.27
Sub G1 50.3

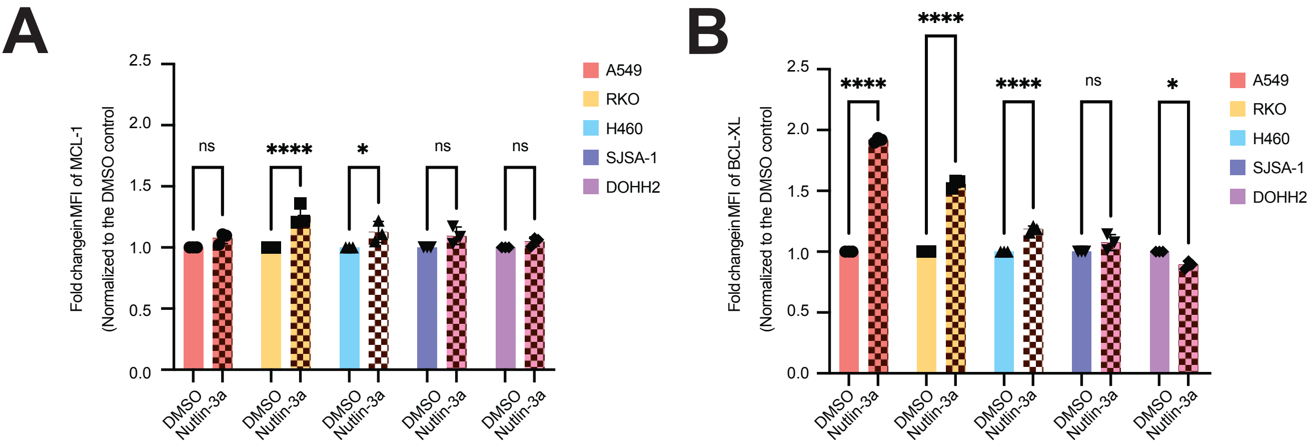
A
Fold change in MFI of MCL-1
(Normalized to the DMSO control
2.5
2.0
1.5
1.0
0.5
0.0
ns
****
*
ns
ns
DMSO
Nutlin-3a
A549
RKO
H460
SJSA-1
DOHH2
B
Fold change in MFI of BCL-XL
(Normalized to the DMSO control
****
****
****
ns
*

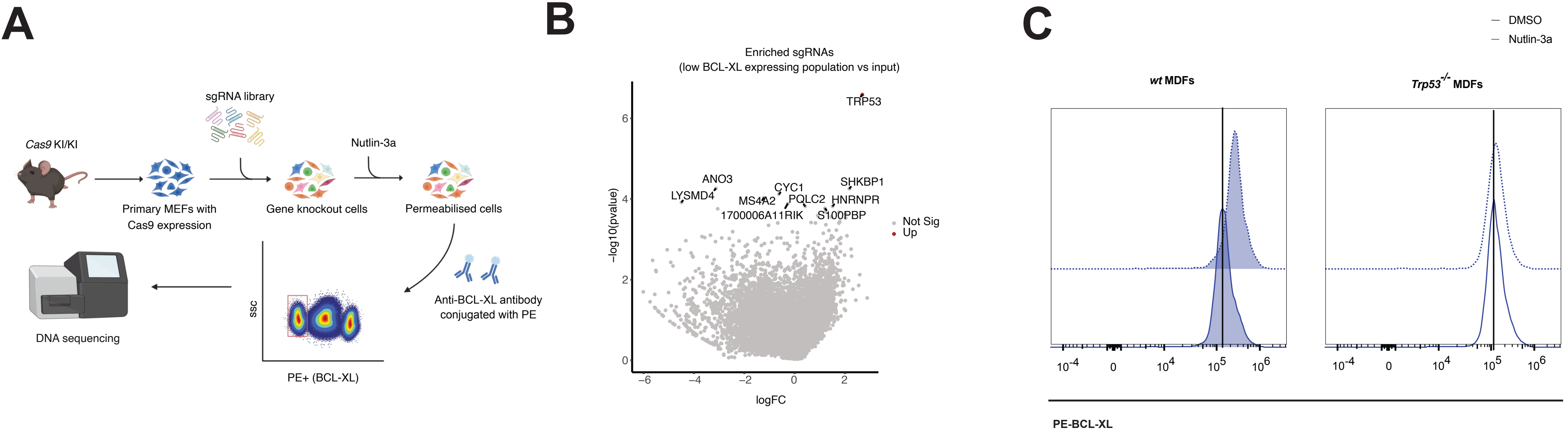
A
sgRNA library
Cas9 KI/KI
Nutlin-3a
Primary MEFs with
Cas9 expression
Gene knockout cells
Permeabilised cells
Anti-BCL-XL antibody
conjugated with PE
DNA sequencing
ssc
PE+ (BCL-XL)
B
Enriched sgRNAs
(low BCL-XL expressing population vs input)
TRP53
ANO3
LYSMD4
CYC1
MS4A2
POLC2
SHKBP1
HNRNPR
1700006A11RIK
S100PBP
Not Sig
Up
−log10(pvalue)
logFC
C
DMSO
Nutlin-3a
wt MDFs
Trp53-/- MDFs
PE-BCL-XL

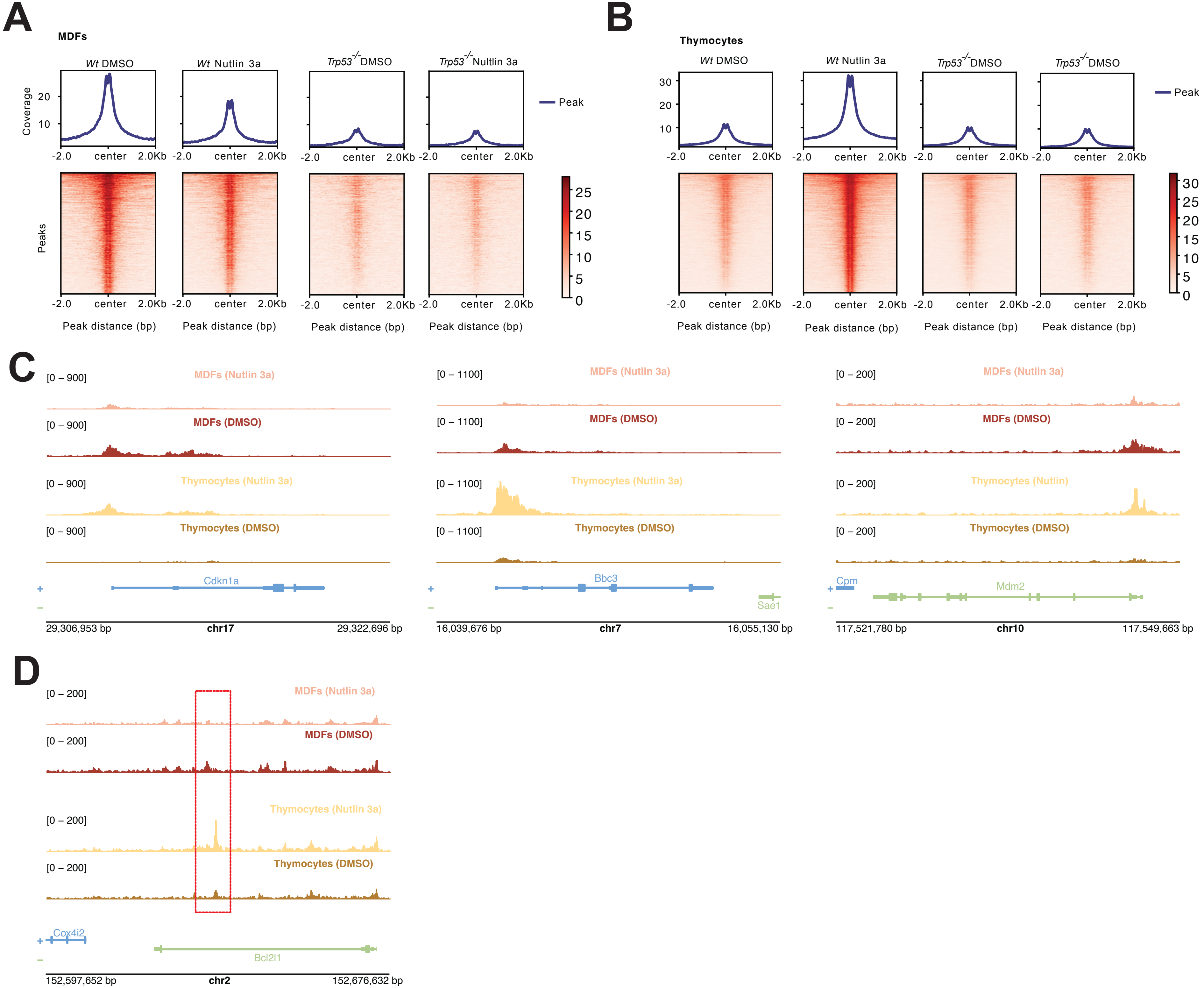
A
MDFs
Wt DMSO
Wt Nutlin 3a
Trp53-/- DMSO
Trp53-/- Nutlin 3a
Coverage
Peak
-2.0 center 2.0Kb
Peaks
Peak distance (bp)
B
Thymocytes
Wt DMSO
Wt Nutlin 3a
Trp53-/- DMSO
Trp53-/- DMSO
Peak
-2.0 center 2.0Kb
Peak distance (bp)
C
[0 – 900]
MDFs (Nutlin 3a)
MDFs (DMSO)
Thymocytes (Nutlin 3a)
Thymocytes (DMSO)
Cdkn1a
29,306,953 bp
chr17
29,322,696 bp
[0 – 1100]
Bbc3
Sae1
16,039,676 bp
chr7
16,055,130 bp
[0 – 200]
Thymocytes (Nutlin)
Cpm
Mdm2
117,521,780 bp
chr10
117,549,663 bp
D
[0 – 200]
Cox4i2
Bcl2l1
152,597,652 bp
chr2
152,676,632 bp
